# Sen1 is a key regulator of transcription-driven conflicts

**DOI:** 10.1101/2022.02.09.479708

**Authors:** Umberto Aiello, Drice Challal, Griselda Wentzinger, Armelle Lengronne, Rowin Appanah, Philippe Pasero, Benoit Palancade, Domenico Libri

**Affiliations:** Université Paris Cité, CNRS, Institut Jacques Monod, F-75013 Paris, France; Institut de Biologie Intégrative de la Cellule UMR9198 CEA, CNRS, Université Paris-Saclay, F-91198 Gif-sur-Yvette, France; Institut de Génétique Humaine, CNRS, Université de Montpellier, Montpellier, France; Genome Damage and Stability Centre, School of Life Sciences, University of Sussex, Falmer, Brighton BN1 9RQ, UK; Institut de Génétique Moléculaire de Montpellier, Univ Montpellier, CNRS, Montpellier, France

**Keywords:** transcription, replication, transcription-replication conflicts (TRCs), Sen1, RNase H, R-loops, genome stability, non-coding transcription, H-CRAC

## Abstract

Cellular homeostasis requires the coordination of several machineries concurrently engaged on the DNA. Wide-spread transcription can interfere with other processes and transcription-replication conflicts (TRCs) threaten genome stability. The conserved Sen1 helicase terminates non-coding transcription, but also interacts with the replisome and reportedly resolves genotoxic R-loops. Sen1 prevents genomic instability but how this relates to its molecular functions remains unclear. We generated high-resolution, genome-wide maps of transcription-dependent conflicts and R-loops using a Sen1 mutant that has lost interaction with the replisome but is termination proficient. We show that Sen1 removes RNA polymerase II at TRCs within genes and the rDNA, but also at sites of transcription-transcription conflicts under physiological conditions, thus qualifying as a “key regulator of conflicts”. We demonstrate that genomic stability is only affected by Sen1 mutation when, in addition to its role at the replisome, termination of non-coding transcription or R-loop removal are additionally compromised.

## INTRODUCTION

The DNA is a shared workspace synchronously used by many cellular machineries that are essential for the correct expression, maintenance, repair, and transmission of the genetic information. The orchestration of these activities must be accurately coordinated, to avoid interferences that might ultimately lead to mis-expression or corruption of the genetic content. RNA Polymerase II (RNAPII) transcribes largely beyond the limits dictated by apparent physiological significance, a phenomenon dubbed pervasive transcription, which has a large impact in genomic crowding. Transcription-replication conflicts (TRCs) can generate genomic instability and jeopardize the faithful transmission of genetic information, in particular when associated to the formation of R-loops. These structures are characterized by a peculiar topological arrangement in which the nascent RNA associates to its DNA template, leaving unpaired the cognate DNA strand. R-loops have important physiological functions in the generation of antibody diversity, and other processes (for a review see: Feng et al., 2020), but their non-physiological accumulation is generally considered genotoxic.

The helicase Sen1 has a particular place in the orchestration of transcription and replication activities. Within the Nrd1-Nab3-Sen1 (NNS) complex, it has an essential role in controlling transcription termination at thousands of genes producing non-coding RNAs (Steinmetz et al., 2006; Hazelbaker et al., 2012; Porrua and Libri, 2013; Schaughency et al., 2014). Failures in NNS-dependent termination has been shown to generate extended transcription events that affect the expression of neighboring genes, thus altering the overall transcriptional homeostasis of the cell (Schulz et al., 2013). Besides a role in limiting the chances of conflicts by restricting pervasive transcription, Sen1 has been proposed to work directly at sites of TRCs. Sen1 loss-of-function mutants, or strains in which the protein has been depleted, display genomic instability phenotypes revealed by increased mitotic recombination between direct repeats, synthetic lethality with DNA repair mutants and Rad52 *foci* accumulation (Mischo et al., 2011). These effects have been attributed to the defective resolution of R-loops at TRCs in the light of increased fork stalling at sites of convergent transcription and replication and the genomic co-localization of Sen1 and replication forks (Alzu et al., 2012). Indeed, increased R-loop levels have been detected in these Sen1 loss-of-function genetic backgrounds especially during S-phase (Mischo et al., 2011; San Martin-Alonso et al., 2021), which led to the proposal that Sen1, by virtue of its helicase activity, resolves R-loops that are formed at TRCs. However, in these mutant contexts, transcription termination of many ncRNA genes is affected, potentially leading to more R-loops formed as a consequence of the generally higher transcriptional readthrough, entailing increased chances of conflicts. Although genomic instability was not observed in other mutants of the NNS complex that have termination defects (Costantino and Koshland, 2018; Mischo et al., 2011), it remains possible that the phenotypes associated to Sen1 loss-of-function originate from the synthetic association of increased transcriptional challenges and defective functions at TRCs . Disentangling the contributions of these potential synthetic effects is paramount for understanding the function of Sen1 in maintaining genomic stability.

We have recently reported the physical interaction of Sen1 with the Ctf4 and Mrc1 replisome components and characterized a mutant, *sen1-3*, that loses this interaction (Appanah et al., 2020). *Sen1-3* cells have a minor growth phenotype and, importantly, no transcription termination defects at NNS target genes. However, this mutation induces lethality in the absence of the two yeast RNases H, Rnh1 and Rnh201/202/203, which, together with other genetic interactions with mutants of the fork stalling signaling pathway (Appanah et al., 2020), underscores the physiological relevance of the interaction of Sen1 with the replisome. Importantly, RNase H1 and H2 are redundantly involved in the degradation of the RNA moiety of R-loops, which might mechanistically underlie a functional connection with Sen1 at TRCs.

In the course of a separate study we have shown that Sen1 is required for a back-up mechanism of RNAPIII release when primary termination has failed (Xie et al., 2021). The interactions of Sen1 with the replisome and RNAPIII are mutually exclusive and sensitive to the *sen1-3* mutation, which implies the existence of two distinct Sen1-containing complexes presumably with different functional roles. These findings indicate that the *sen1-3* mutant allows untangling the function of Sen1 in NNS-dependent termination from its functions at the replisome and RNAPIII transcription.

Here we first addressed the functional impact of the *sen1-3* mutation on transcription-replication conflicts under conditions in which neither transcription nor replication are altered. Prompted by the strong genetic interaction with RNases H we also studied the role of these enzymes at sites of conflicts and the impact of R-loops. To this aim we generated high-resolution transcription maps in different phases of the cell cycle in *sen1-3* cells and in the absence of RNases H. We also devised a novel methodology to detect R-loops *in vivo* with high sensitivity and unprecedented resolution. We show that Sen1 is required for the efficient removal of RNAPII at many TRC sites within genes, but also at the rDNA, where it collaborates with RNases H. Surprisingly, when non-coding transcription is correctly terminated, loss of the interaction of Sen1 with the replisome does not cause increased R-loop accumulation, mitotic recombination or DNA damage as observed in *sen1* loss-of-function mutants. We demonstrate that increased DNA damage observed in these mutants requires both the lack of Sen1 interaction with the replisome and the transcription termination defects at non-coding RNA genes. Interestingly, we show that Sen1 also functions at many other genomic sites to remove RNAPII at sites of conflicts with RNAPIII and possibly RNAPI. We propose a model according to which Sen1 has key functions in the regulation of transcription-driven conflicts in the genome, and we demonstrate that these functions are independent from its role in terminating non-coding transcription.

## RESULTS

### Sen1 depletion leads to major genome-wide alterations in the transcriptome

To assess directly the possible contribution of transcription termination defects to the genomic instability phenotypes observed in Sen1 loss-of-function mutants, we first directly gauged the extent of alterations in coding and non-coding RNAPII transcriptional activity under defective Sen1 function. We generated high resolution transcription maps using RNAPII CRAC (Crosslinking Analysis of cDNAs, Bohnsack et al., 2012; Candelli et al., 2018) upon depletion of Sen1 by the auxin degron system (Nishimura et al., 2009). The CRAC methodology allows detecting the position of RNAPII with directionality and high resolution.

Depletion of Sen1 by addition of auxin induced the expected transcription termination defects at canonical NNS targets (CUT and snoRNA genes, Figures S1A, left panel, and S1B), but not at mRNA coding genes (Figure S1A, right panel), consistent with a previous study (Schaughency et al., 2014). To estimate the occurrence of global, extragenic transcriptional changes in an unbiased manner, we computed the RNAPII CRAC signal in 200 nt, non-overlapping windows with the exclusion of mRNA-coding genes, the rDNA and tRNA genes to first focus on regions of direct Sen1 action. Scatter plots of these values revealed a dramatic alteration of intergenic transcription upon Sen1 depletion compared to wild-type cells (Figure 1A, compare left and right). Similar transcription alterations were observed upon auxin-dependent depletion of another NNS component, Nrd1, indicating that they are linked to a transcription termination defect (Figure S1C).

**Figure 1:**
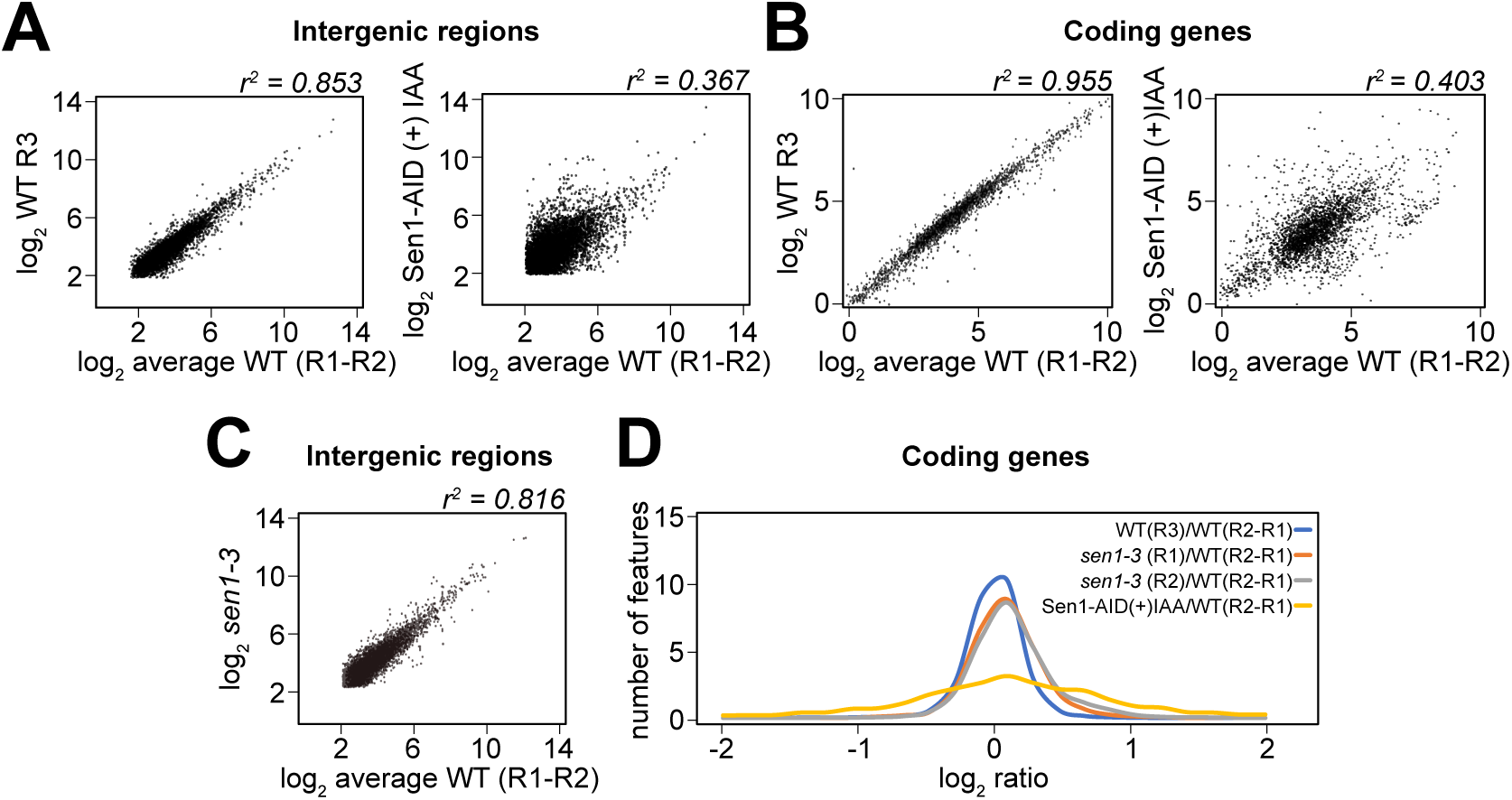
Major alterations in the transcriptome upon Sen1 depletion. **A)** Scatter plots of RNAPII CRAC log_2_ values computed in non-overlapping 200 nt bins relative to the average of two wild-type (WT) replicates used as a common reference in all panels. Right: Sen1-AID strain under depletion conditions (auxin added for 1 hour); left: another WT replicate for comparison. mRNA-coding and tRNA genes have been excluded from the calculations. Only the W strand has been analysed, which also excludes the rDNA *locus*. Bins containing low number of reads (log_2_<2) have been excluded from the analysis for clarity. **B)** As in A but RNAPII CRAC signal has been computed on mRNA-coding genes. **C)** As in A, right panel, intergenic values of one replicate from *sen1-3* cells is compared to the WT reference. **D)** Distributions of log_2_ ratios of average RNAPII CRAC signals detected on mRNA-coding genes. Data from two independent *sen1-3* replicates, evaluated relative to the WT reference. An additional WT replicate is shown for comparison as well as the distribution obtained upon auxin depletion of Sen1-AID.

The gene expression program was also significantly modified upon Sen1 depletion (Figure 1B), with many stress genes activated and ribosomal protein genes downregulated. These effects were generally recapitulated upon depletion of Nrd1, the two datasets showing a high level of correlation (r^2^=0.726, Figure S1D). Because we did not observe termination defects at genes with altered expression (and at mRNA genes in general, Figure S1B, first panel), these effects are likely due to a cellular stress response induced by Sen1 (and Nrd1) depletion.

These results demonstrate that loss of full Sen1 function has a major impact on the distribution of transcription events genome-wide and on gene expression. These changes are expected to alter the wild-type landscape of TRCs and might contribute significantly to the genomic instability phenotypes observed in Sen1 loss-of-function mutants.

### Interaction of Sen1 with the replisome is required to solve TRCs in the 5’-end of genes

In the light of the above considerations, we sought to analyze the role of Sen1 in genomic stability in a more physiological transcriptional landscape. Genetic data strongly support the notion that the role of Sen1 at TRCs is mechanistically linked to its interaction with the replisome, and therefore we turned to the use of the *sen1-3* mutant that loses this interaction, but is proficient for NNS termination (Appanah et al., 2020). In sharp contrast to what observed upon depletion of Sen1, affecting the interaction of Sen1 with the replisome does not alter significantly the coding and non-coding transcriptomes (Figures 1C and 1D), validating our strategy.

Failure to resolve or to avoid a TRC is expected to result in slowing down or stalling of a replication fork but also to induce the accumulation of RNAPII at the site of conflict. We focused on the transcription side of the conflict and reasoned that if Sen1 is recruited at the replisome to terminate conflicting transcription, in its absence RNAPII should accumulate at these sites, thus providing a signature of Sen1-dependent conflicts.

Assuming that conflicts depend, at least to some extent, on the stochastic encounters of the two machineries, they can be expected to be more frequent in genomic regions with inherently higher RNAPII occupancy. Higher RNAPII persistency is observed in the 5’ region of yeast genes where the elongation complex is known to pause (Figure S1A, right panel; see also Churchman and Weissman, 2011; Mayer et al., 2011). Note that 5’-end proximal pausing is an inherent feature of the transcription process that is independent of replication as it is observed also in cells arrested in G_1_ (see below, Figure 2D).

**Figure 2:**
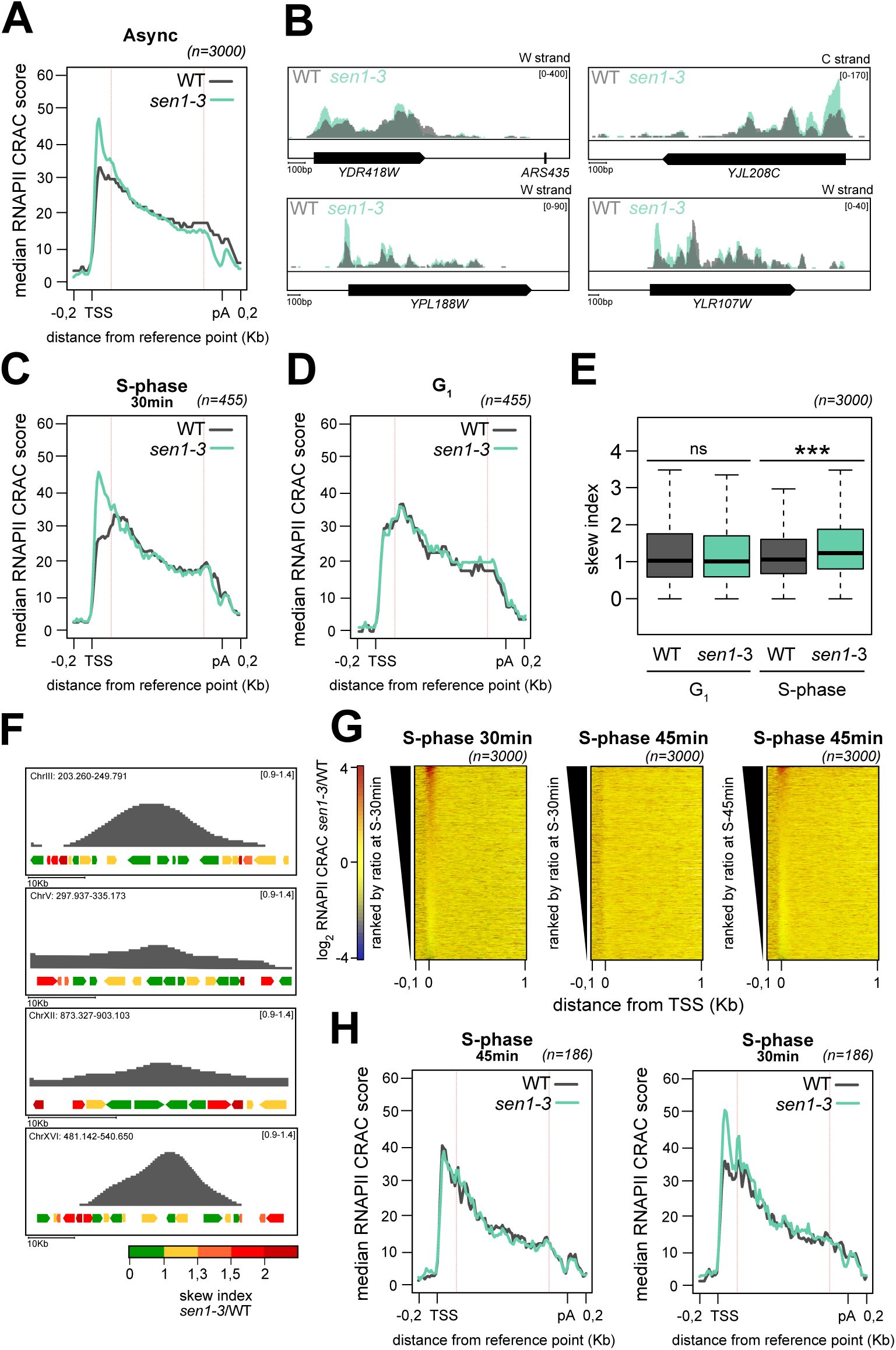
Sen1 promotes release of RNAPII from the 5’-end of genes during replication. **A)** Metagene analysis of the RNAPII distribution at mRNA-coding genes aligned at their TSS and at their pA site in WT and *sen1-3* cells grown asynchronously. The 3000 genes with the highest expression were used for the analysis. Values on the y-axis correspond to the median coverage. Only the region in between the red lines is scaled. **B)** Integrative Genomics Viewer (IGV) representative screenshots of genes displaying RNAPII accumulation in *sen1-3* cells. The overlap of RNAPII read coverage in WT (grey) as and *sen1-3* cells (aqua green) is shown. The strand (W for Watson, C for Crick) and a scale bare are indicated respectively in the top-right and the bottom-left corners. **C)** Metagene analysis as in A, but for the genes with the highest skew index *sen1-3*/WT ratio in cells synchronously released in S-phase for 30 min. **D)** Metagene analysis as in C but for cells arrested in G_1_. **E)** Comparison of the gene skew ratio for the indicated strains in G_1_ or in S-phase (30 min). *** p<0.001**. F)** Examples of replicons as detected by DNA copy number analyses for cells synchronously released in S-phase for 30 min (signal from *sen1-3* cells). The genomic coordinates are indicated at the top left of each example. Genes were color-coded based on their skew index ratio as indicated. **G)** Heatmaps representing the RNAPII CRAC signal change (log_2_ ratio) in the *sen1-3* mutant relative to the WT for mRNA coding genes aligned at their TSS in S-phase for 30 min and 45 min as indicated. Genes were ranked according to the signals detected in the first 200 nt after the TSS at the 30 min (left and central panel) or 45 min time point (right panel). **H)** Left: Metagene analysis as in Figure 2C, but on cells released in S-phase for 45 minutes and on a group of genes already replicated at this time point. Right: the same group of genes as in Left, but at the earlier 30 min time point.

We therefore first analysed Sen1-dependent alterations on RNAPII distribution focusing on the regions of 5’-proximal transcriptional pausing. Marked changes in the RNAPII profile was clearly observed by metasite analyses and by the inspection of individual genes in asynchronously growing *sen1-3* cells, with an increase in occupancy specifically in the 5’-end (Figures 2A and 2B). In some cases, increased occupancy was also observed at other sites of RNAPII pausing (e.g., see *YDR418W* in Figure 2B). This pattern is not compatible with increased transcription initiation at a set of genes, which would result in a homogeneous, increase of the RNAPII CRAC signal. Rather, it points to the occurrence of increased RNAPII pausing, mainly in the 5’-end of genes, possibly due to defective RNAPII release at sites of conflicts when Sen1 is absent from the replisome.

To assess the dependency on ongoing replication, RNAPII CRAC was performed using cells arrested in G_1_ by the addition of alpha factor and synchronously released in S-phase. The progression of replication was analysed in the same cells by DNA copy number analyses. At 30 minutes after the release in S-phase, 5’-skewed RNAPII occupancy was observed in a subset of genes in *sen1-3* cells (Figure S2A; see also Figure 2G), recapitulating what observed in asynchronous cells. For a quantitative analysis of these observations, we calculated a skew index, defined as the ratio of the RNAPII CRAC score in the 5’-end [TSS; TSS+200] to the signal in a downstream region of identical length [TSS+300; TSS+500]. The skew index is expected to provide a readout that is independent from changes in transcription initiation levels, which might occur stochastically or between WT and *sen1-3* cells. For more robust analyses, we also focused on the subset of most affected genes, identified by computing the skew index ratio (*sen1-3/*WT) for each gene and selecting features one standard deviation over the mean. This resulted in a set of 455 genes, with a marked RNAPII CRAC signal increase in *sen1-3* cells, positioned slightly upstream of the canonical 5’ peak detected in the wild-type (Figure 2C). Most importantly, 5’ skewed RNAPII occupancy was not observed in the absence of replication, when cells were arrested in G_1_ (compare Figures 2C and 2D) and the distributions of skew indexes were significantly different only in S-phase (Figure 2E). This conclusion also holds genome-wide, when considering the set of most expressed 3000 genes (Figure S2B). These findings indicate that replication is required for the increased occupancy of RNAPII observed in the 5’-end of genes in *sen1-3* cells, supporting the notion that RNAPII is not released efficiently at TRCs when Sen1 cannot interact with the replisome.

If the replication-dependent RNAPII increase in the 5’-end of genes is linked to TRCs, the group of affected genes should be located in regions where replication is ongoing. To address this point, we color-coded genes based on their skew index and monitored their distribution along the chromosomes in relation to the position of the replicative forks as detected by DNA copy number analysis. The normalized DNA level signal in replication regions (Figures 2F and S2C, see Materials and Methods) is linked to the probability that a given sequence has been replicated. Regions with the highest chance of having been replicated are located in the center of the replicon, while sequences undergoing replication are located close to its periphery. Genes with the highest skew index ratio were not randomly distributed in the active replication region but preferentially positioned around the borders (Figures 2F, S2C-D). Symmetrically, genes positioned at the borders have a statistically significant higher skew ratio than genes in the center of the replication region (Figure S2E). These data are fully compatible with the notion that genes with a high skew ratio are undergoing replication.

Genes in a co-directional (CD) and head-on (HO) orientation relative to the direction of replication were equally found to be affected (Figures 2F, S2C and S2F), as also shown by measuring the distance of the closest origins generating HO or CD replication for each affected gene (Figure S2G). This suggests that Sen1 can solve both kinds of conflicts by binding to the replisome.

One prediction of our model is that the set of genes undergoing TRCs should change as replication progresses. To test this prediction, we generated additional RNAPII CRAC transcription maps at 45 min after release in S-phase and compared the results to the transcription maps that were generated at the earlier 30 min time point. The RNAPII CRAC signal generated at a later replication time point recapitulated the phenotype observed at the 30 min time point (Figure S2H and 2G, right panel), yielding a group of 439 affected genes (Figure S2J) selected following the aforementioned criteria. However, when the genes ranked for increased RNAPII occupancy at the 30 min time point were monitored for RNAPII CRAC signals at the 45 min time point (Figure 2G, compare left and middle panel), a very poor overlap, if any, was observed, consistent with the notion that TRCs are restricted to the set of genes undergoing replication at a given time point. Consistent with this notion, 186 genes selected for having already been replicated at the 45 min time point did not show the 5’-skewed RNAPII signal in *sen1-3* cells (Figure 2H, left panel).

In agreement with the presence of a challenged replication environment, the size of early replicons was found to be significantly smaller in *sen1-3* cells relative to the WT (Figure S2K, left panel). Interestingly, the same analysis also revealed the premature activation of late origins in the *sen1-3* mutant (Figure S2K, right panel), explaining the similar length of S-phase observed in *sen1-3* and WT cells (Appanah et al., 2020).

Together, these findings indicate that ongoing replication is required for the accumulation of RNAPII in the 5’-end of a set of genes, which we propose to be due to the failure to remove RNAPII from sites of TRCs when Sen1 cannot interact with the replisome. They also suggest that TRCs are frequently occurring even under physiological conditions and that the failure to efficiently prevent or resolve them alters the replication program.

### Association of Sen1 with the replisome is required for limiting RNAPII accumulation at the ribosomal replication fork barrier in S-phase

The ribosomal DNA Replication Fork Barrier (rRFB) is a site where one replication fork stalls upstream of the DNA-bound Fob1 protein at the 3’-end of ribosomal DNA repeats (Kobayashi, 2003). This ensures that each rDNA repeat is being replicated in a co-directional fashion with RNAPI and RNAPIII transcription (Figure 3A).

**Figure 3:**
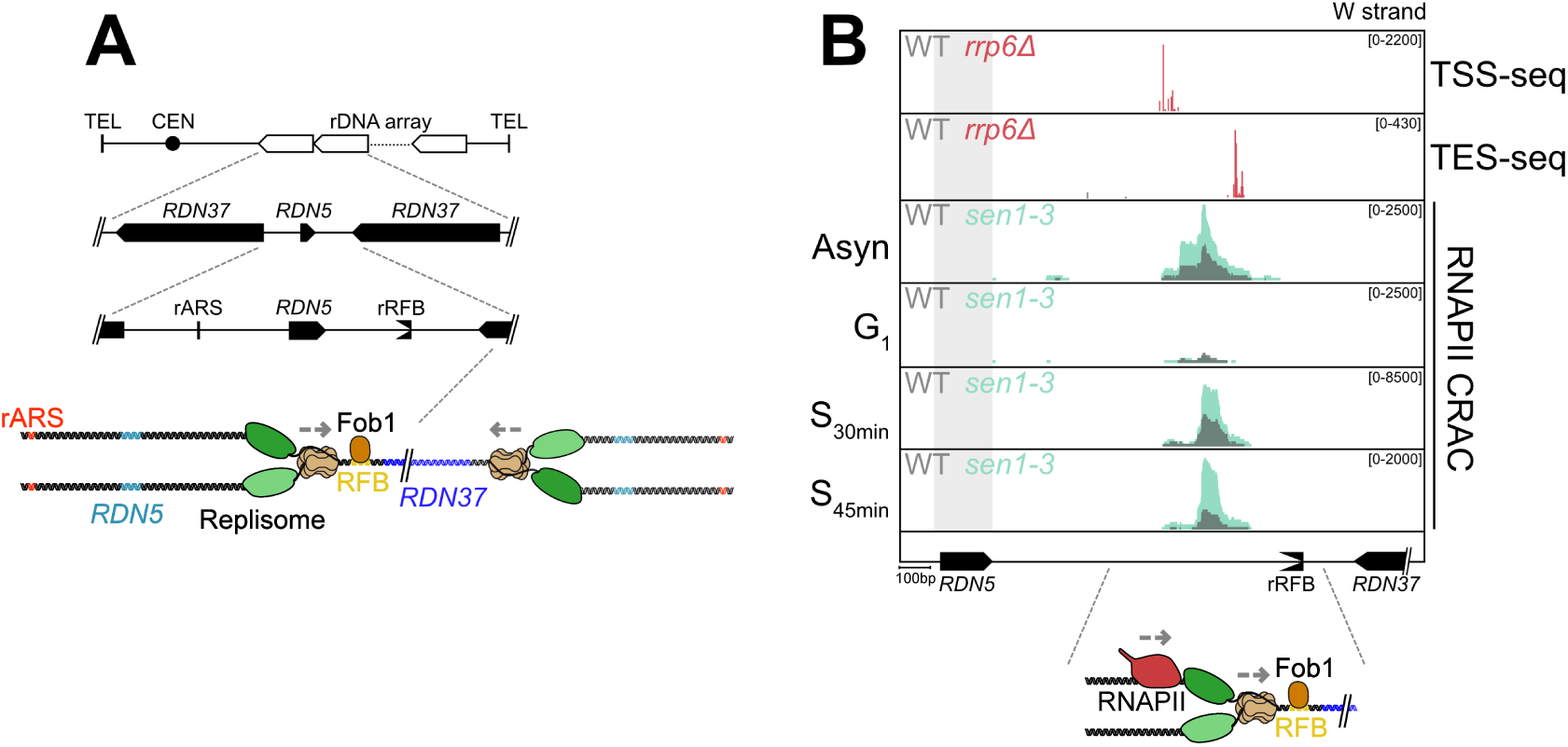
Association of Sen1 with the replisome is required for limiting TRCs at the ribosomal replication fork barrier. **A)** Schematic representation of the rDNA *locus* on ChrXII. The position of the replication origin (rARS) and of the rRFB are indicated relative to the *RDN37* and *RDN5* genes. A cartoon illustrates the direction of replication in the region (grey dashed arrow) and the function of the Fob1 protein. **B)** Screenshots illustrating the distribution of the RNAPII CRAC signal density around the rRFB in the strains and the conditions indicated as in Figure 2. The TSS and the TES of the unstable RNA produced by the fork-trailing transcription event are shown in the first and second tracks (Challal et al., 2018a). At the bottom of the panel, a cartoon illustrates the presumed position of the stalled replisome and RNAPII.

Monitoring RNAPII by CRAC in the rRFB region in asynchronous cells revealed the existence of a non-annotated transcription unit located upstream and in close proximity of the rRFB, which generates a cryptic unstable transcript that can only be detected in an exosome-defective, *rrp6*Δ background (Figure 3B). Transcription in this region might generate a co-directional conflict with forks stalled at the rRFB and be of interest for our analysis.

Interestingly, the RNAPII CRAC signal was close to background in G_1_-arrested cells yet was markedly visible both at the earlier and later replication time points (Figure 3B), indicating that accumulation of RNAPII only occurs during or after replication, possibly as a consequence of fork stalling at the Fob1-bound RFB. Consistent with this notion, the RNAPII peak is located roughly 100 nt from the edge of the rRFB, which is hardly reconcilable with Fob1 roadblocking RNAPII (Candelli et al., 2018; Colin et al., 2014) but fully compatible with one co-directional replication fork derived from the closest ARS filling the gap between the RNA polymerase and the Fob1-bound rRFB (see scheme in Figure 3B). RNAPII accumulation was RFB-dependent because conditional depletion of Fob1 by the auxin degron system (Nishimura et al., 2009), resulted in a significant decrease in the RNAPII signal as detected by Chromatin Immunoprecipitation (ChIP) (Figure S2M). Most importantly, the RNAPII CRAC signal was found to be considerably increased in *sen1-3* cells indicating that the interaction of Sen1 with the replisome is required for releasing RNAPII at this site while in close proximity with a replication fork. Increased RNAPII signal in *sen1-3* cells was not due to increased rDNA copy number in this strain as verified by qPCR (see below, Figure 6D). Analysis of published ChIP-exo data (Rossi et al., 2021) confirmed the specific presence of Sen1 at the rDNA in close correspondence with the RNAPII CRAC peak (Figure S4A).

Together these data provide evidence for replication-dependent RNAPII accumulation, most likely in the wake of the replication fork stalled at the rRFB. Importantly, they also demonstrate that the interaction of Sen1 with the replisome is required for the efficient release of RNAPII paused in close proximity with the replisome.

### R-loops are detected at the rRFB by H-CRAC

Several studies have linked mutation or depletion of Sen1 to the accumulation of R-loops at TRCs. The requirement of RNase H activity for the viability of *sen1-3* cells suggests that degradation of R-loops is essential in at least some genomic locations when Sen1 cannot interact with the replisome. Therefore, we decided to generate genome-wide maps of R-loops taking advantage of the unprecedented benefits offered by the *sen1-3* mutant.

Currently available tools to produce genome-wide R-loops maps (R-ChIP, DRIP-seq and related techniques) often lack directionally, do not always provide consistent outputs and have a limited resolution (Chédin et al., 2021), which we feared to not be sufficient for an integration with our RNAPII CRAC data. To overcome these limitations, we reasoned that since RNase H binds and degrades the RNA moiety of DNA:RNA heteroduplexes, it should be possible to catch it in action at its targets *in vivo* by UV-crosslinking. Sequencing of the associated RNA would thus provide a sensitive and high-resolution detection of DNA:RNA hybrids/ R-loops.

H-CRAC (for RNase <u>H</u> - <u>CRAC</u>) experiments performed with both RNase H1 and RNase H2 provided very reproducible and similar, stand-specific outputs, despite revealing some specificities (Figure 4A and S3A, see below). This was expected in the light of the known redundancy of these enzymes in R-loop degradation.

**Figure 4:**
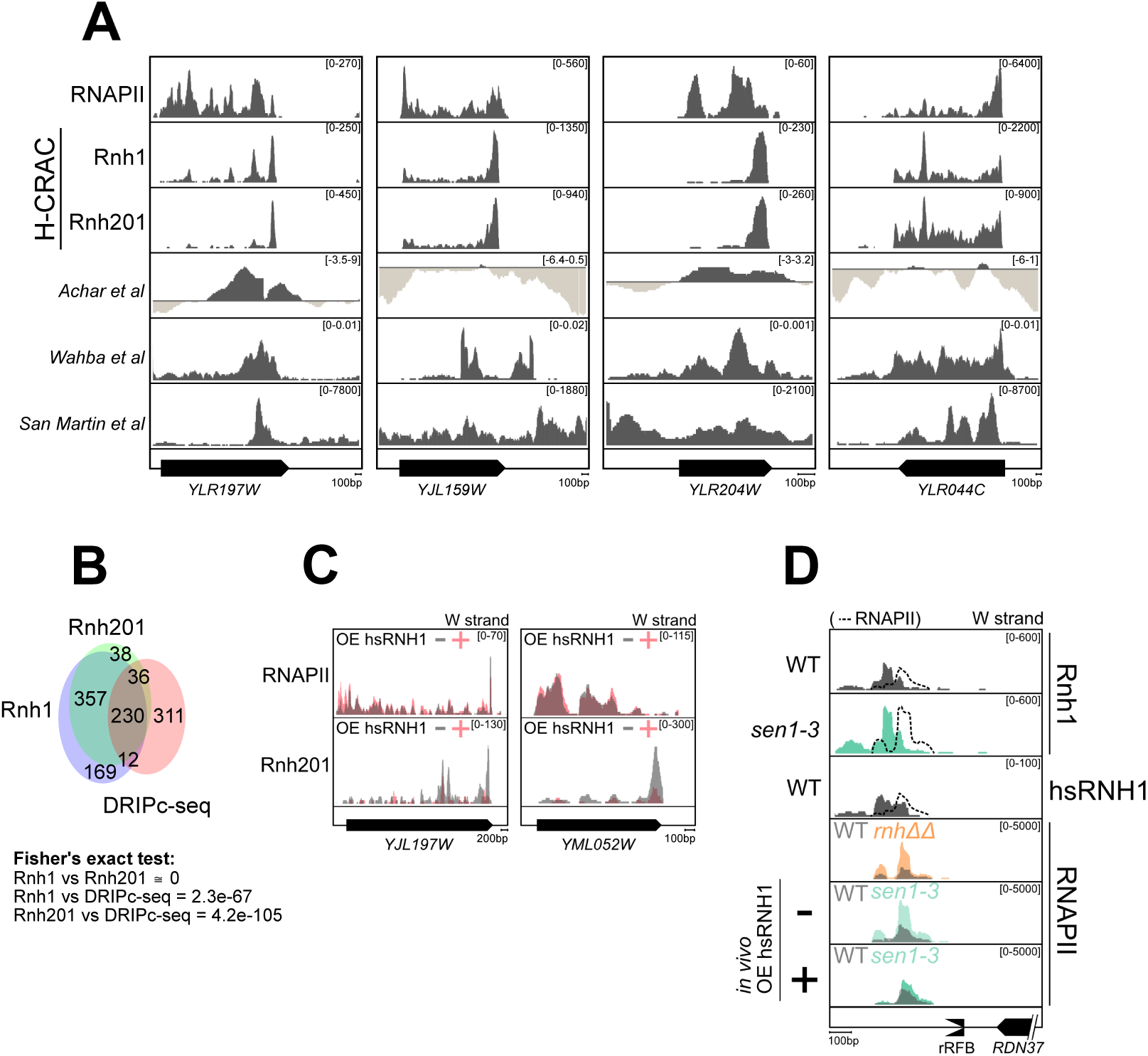
Roles of R-loops and RNase H in Sen1-dependent RNAPII release at the RFB. **A)** Comparison between DRIP-seq (Achar et al., 2020; San Martin-Alonso et al., 2021; Wahba et al., 2016) and H-CRAC signals detected at individual genes. The RNAPII CRAC track is shown for evaluating the R-loop signals relative to transcription. For the directional DRIP-seq (San Martin-Alonso et al., 2021), the H-CRAC and the RNAPII CRAC only the strand of the target gene is shown. **B)** Overlap of R-loop prone genes (HTseq > mean +1sd) defined by the directional DRIP-seq (San Martin-Alonso et al., 2021) and by H-CRAC (Rnh1 or Rnh201). The significance of the overlap between H-CRAC and the DRIP-seq dataset was calculated with Fisher’s exact tests and is indicated at the bottom. **C)** Examples of decrease in H-CRAC (Rnh201) signals upon *in vivo* overexpression of hsRNH1, despite no effects on transcription as shown by RNAPII CRAC tracks. **D)** R-loops and RNAPII accumulation at the rRFB as detected by Rnh1, Rnh201 and hsRNH1 H-CRAC and RNAPII CRAC. The position where RNAPII accumulates in the corresponding background is indicated by a dashed curve. Increased RNAPII occupancy is observed at the rRFB in *rnh1Δ rnh201Δ* (*rnhΔΔ*) cells. RNAPII increased occupancy is suppressed by *in vivo* overexpression of hsRNH1.

We gauged the validity of the proposed landmarks for R-loop detection (Chédin et al., 2021) by first ectopically expressing in yeast a sequence derived from the mouse AIRN gene that was demonstrated to form R-loops *in vitro* and *in vivo* (Carrasco-Salas et al., 2019; Ginno et al., 2012). We verified by DRIP that this sequence generates high levels of RNase H-sensitive R-loops when transcribed in *S. cerevisiae* (Figure S3B). Prominent H-CRAC signals were detected at the ectopically expressed mAIRN *locus* (Figure S3C), validating the notion that H-CRAC robustly identifies well-established regions of R-loop formation.

H-CRAC signals overlapped transcription at the genome-wide scale, as expected considering the co-transcriptional nature of R-loop formation (Figures 4A, S3D and S3E), but the pattern of the RNAPII and RNase H signals was generally different (Figures 4A and S3E), consistent with the notion that not all transcribed regions generate R-loops to the same extent. A comparison with the only directional R-loop map generated by S9.6-DRIP-seq in yeast (San Martin-Alonso et al., 2021) revealed significant overlaps when taking into account the overall signal along coding genes (Figure 4B). However, the resolution of the signal was clearly higher and its distribution within genes often different, with prominent H-CRAC signals observed at new locations (Figure 4A and data not shown). The detailed genome-wide analysis of R-loops distribution determined by H-CRAC is beyond the scope of this report and will be provided in a separate manuscript (Aiello et al., in preparation).

Overexpression of human RNH1 considerably reduced Rnh201 H-CRAC signals in many locations (Figure 4C), without a significant, general effect on transcription (Figure 4C and S3F), which was monitored in parallel to ascertain that reduced H-CRAC signals were not due to altered gene expression. We could also generate similar H-CRAC signal distributions using tagged hsRNH1, confirming that the ectopically expressed, heterologous enzyme recognizes very similar targets as the yeast proteins (Figures S3G and S3H). Together, these data demonstrate that H-CRAC is a sensitive and resolutive method for detecting *in vivo* at least a significant fraction of cellular DNA:RNA hybrids, overlapping and complementing *in vitro* DRIP-based methods for detecting R-loops.

To assess whether R-loops form at the rRFB during replication, we generated H-CRAC maps in *sen1-3* and WT cells. Interestingly, we detected prominent Rnh1 H-CRAC signals peaking roughly 50 nt upstream of the fork-trailing RNAPII peak (Figures 3B and 4D), consistent with the notion that they form immediately upstream of the stalled RNA polymerase. Rnh201 signals were less prominent in this position and were instead preferentially observed roughly 500 nt upstream (Figure S4A), suggesting that Rnh1 might preferentially recognize R-loops at this site. Ectopically-expressed human RNase H (hsRNH1) for H-CRAC generated clear signals that nicely overlapped yeast Rnh1 targets in this region (Figure 4D), further supporting the notion that they represent *bona fide* R-loops.

H-CRAC signals proximal to the rRFB increased significantly in *sen1-3* cells in their absolute levels but not when evaluated relative to the levels of paused RNAPII upstream of the replication fork (Figure 4D). This finding indicates that higher R-loop levels at this site parallel the increased stalling of RNAPII engaged in conflicts with replication forks when Sen1 cannot interact with the replisome.

From these experiments we conclude that R-loops form upstream of RNAPII conflicting with the stalled replication forks. The interaction of Sen1 with the replisome is required for releasing fork-trailing RNA polymerases, but not for limiting the levels of R-loops that form *per* transcriptional event.

### RNases H promote RNAPII release at the rRFB

The strong growth defect of *sen1-3* cells in the absence of RNase H activity, suggests that Sen1 at the replisome and RNases H have a common or complementary function either in limiting R-loops accumulation, in RNAPII release, or both. Because we did not observe increased R-loops *per* transcription event at the rRFB in *sen1-3* cells, we considered a possible implication of RNases H/R-loops in RNAPII release. We first monitored RNAPII occupancy by CRAC in a *rnh1Δ rnh201Δ* mutant, which lacks RNase H activity. In this genetic context, transcription was not generally altered, as shown by the profiles of median RNAPII CRAC signals on genes (Figure S4B). Interestingly, however, we observed a clear increase in the levels of RNAPII at the rRFB compared to a WT strain, which accumulated in the same position and to similar levels as in *sen1-3* cells (Figure 4D). Thus, RNases H are required to limit accumulation of RNAPII at the TRC in the rRFB.

RNases H might contribute to the release of the polymerase independently of R-loop degradation, or by degrading the RNA moiety of these structures. In this latter perspective the expected increase in R-loops in *rnh1Δ rnh201Δ*, might prevent efficient RNAPII release. One important prediction of this hypothesis is that degrading the R-loops formed upstream of the stalled polymerase in *sen1-3* cells should favour its release and suppress the RNAPII accumulation phenotype.

Overexpression of human hsRNH1 significantly reduced R-loop levels genome-wide, without generally altering transcription. However, we observed a significant reduction in RNAPII accumulation at the rRFB in *sen1-3* cells (Figure 4D). Together, these results suggest that modulating the levels of R-loops at the rRFB, either by increasing them in *rnh1Δ rnh201Δ* cells or by decreasing them upon overexpression of hsRNH1, affects RNAPII release in an anti-correlative manner. In the light of these results, we considered that RNases H might also contribute to the release of RNAPII at TRCs in the 5’-end of genes undergoing replication. However, the double *rnh1Δ rnh201Δ* mutant did not phenocopy the *sen1-3* RNAPII 5’-end accumulation suggesting that RNases H do not partake in releasing RNAPII in these locations (Figure S4B). This conclusion is also supported by the findings that overexpression of hsRNH1 in *sen1-*3 cells did not suppress 5’-end RNAPII accumulation (Figure S4C, compare with Figure 2A) and that R-loops are not significantly detected in the very 5’-end of genes where the increased RNAPII accumulation is observed (Figures S4D and S4E). One likely explanation for these results is that in the proximity of the TSS the short length of the available nascent RNA is not compatible with the formation of DNA:RNA hybrids (see Discussion).

From these experiments we conclude that RNAPII is released at the rRFB by the combined action of Sen1 at the replisome and RNases H, the latter presumably functioning by degrading the R-loops formed in the wake of the stalled transcription elongation complex.

### Sen1 releases RNAPII at sites of conflicts with RNAPIII

Aside from RNAPII and replisome components, Sen1 also interacts with RNAPIII, an interaction that is also lost *in sen1-3* cells, and is not mediated by the replisome but reflects the existence of an alternative complex (Xie et al., 2021).

We have previously shown that non-coding and non-annotated RNAPII transcription events might collide with RNAPIII transcription units, where they are restrained by roadblocks located in the 5’- and 3’-end of tRNA genes (Candelli et al., 2018). The mechanism by which RNAPII is released at these sites was not addressed, although we showed that RNAPII ubiquitylation and degradation occurs at other sites of roadblock (Colin et al., 2014). We hypothesized that, in analogy with the mechanism described at the replisome, Sen1 might be recruited by RNAPIII for removing conflicting RNAPIIs. We monitored transcription around tRNA genes in WT and *sen1-3* cells and found increased accumulation of RNAPII upstream of tRNA genes in *sen1-3* cells, consistent with our working hypothesis. This can be clearly appreciated at individual cases (Figure 5A) and is statistically significant at the genome-wide scale (Figure 5B, Figure S5A, top panel). We considered the possibility that this accumulation was related to ongoing replication, but the effect proved to be cell cycle independent (Figures 5A, 5B and S5A). Increased accumulation of RNAPII was also observed when monitoring antisense transcription relative to tRNA genes (Figures 5A and 5B, right panels, and S5A, bottom panel), although to lower levels, possibly because additional mechanisms are in place for limiting head-on conflicts. Nevertheless, we clearly observed that, when uncontrolled by Sen1, RNAPII entered tRNA transcription units, at least in the antisense direction (for technical reasons only antisense transcription can be monitored in the body of tRNA genes). Similar to what observed at mRNA coding genes, no significant effect was observed after deletion of the two RNase H genes (Figure S5B), suggesting that degradation of R-loops is not contributing to RNAPII release at these sites.

**Figure 5:**
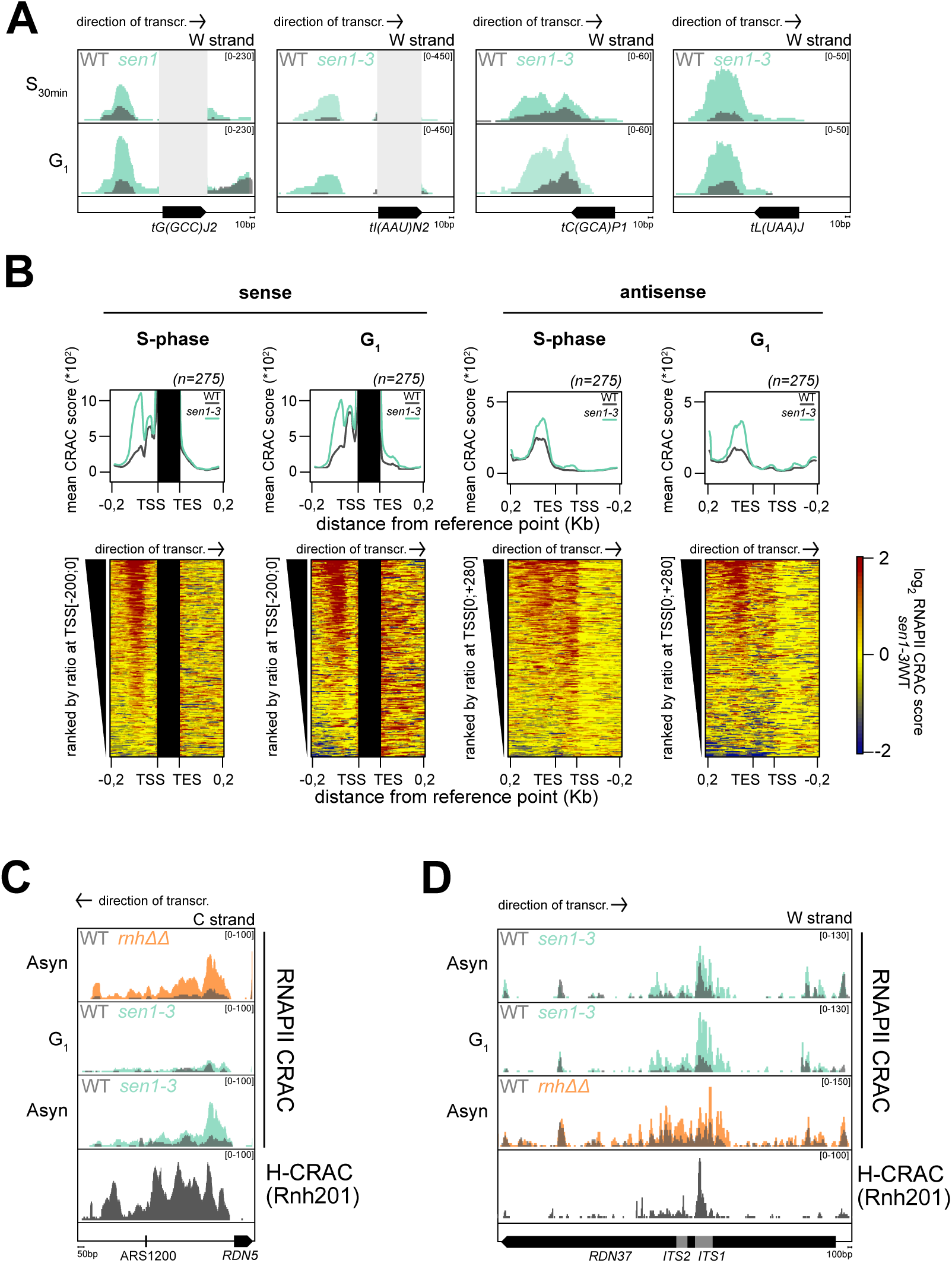
Sen1 releases RNAPII at sites of conflicts with RNAPIII and limits RNAPII transcription in the rDNA. **A)** Snapshots illustrating the RNAPII accumulation in the sense (left panels) or antisense (right panels) orientation at individual tRNA genes in *sen1-3* cells in G_1_ or S-phase (30min) as indicated. Sense signals in the tRNA body are masked as they do not represent *bona fide* RNAPII occupancy but result from contaminating mature tRNAs. **B)** Heatmaps and summary plots representing the log_2_ ratio of RNAPII CRAC signals in *sen1-3* vs WT cells at tRNA genes both for the sense (left) and the antisense transcription (right). Ranking was performed as indicated on the left of each heatmap. As in A, signals within the tRNA body are masked. **C)** RNAPII and R-loop (Rnh201 H-CRAC) densities antisense and upstream of the *RDN5* gene in the indicated mutants. The C strand is shown (antisense of *RDN5*). **D)** As in C but transcription and R-loops antisense to the *RDN37* (W strand) are shown.

**Figure 6:**
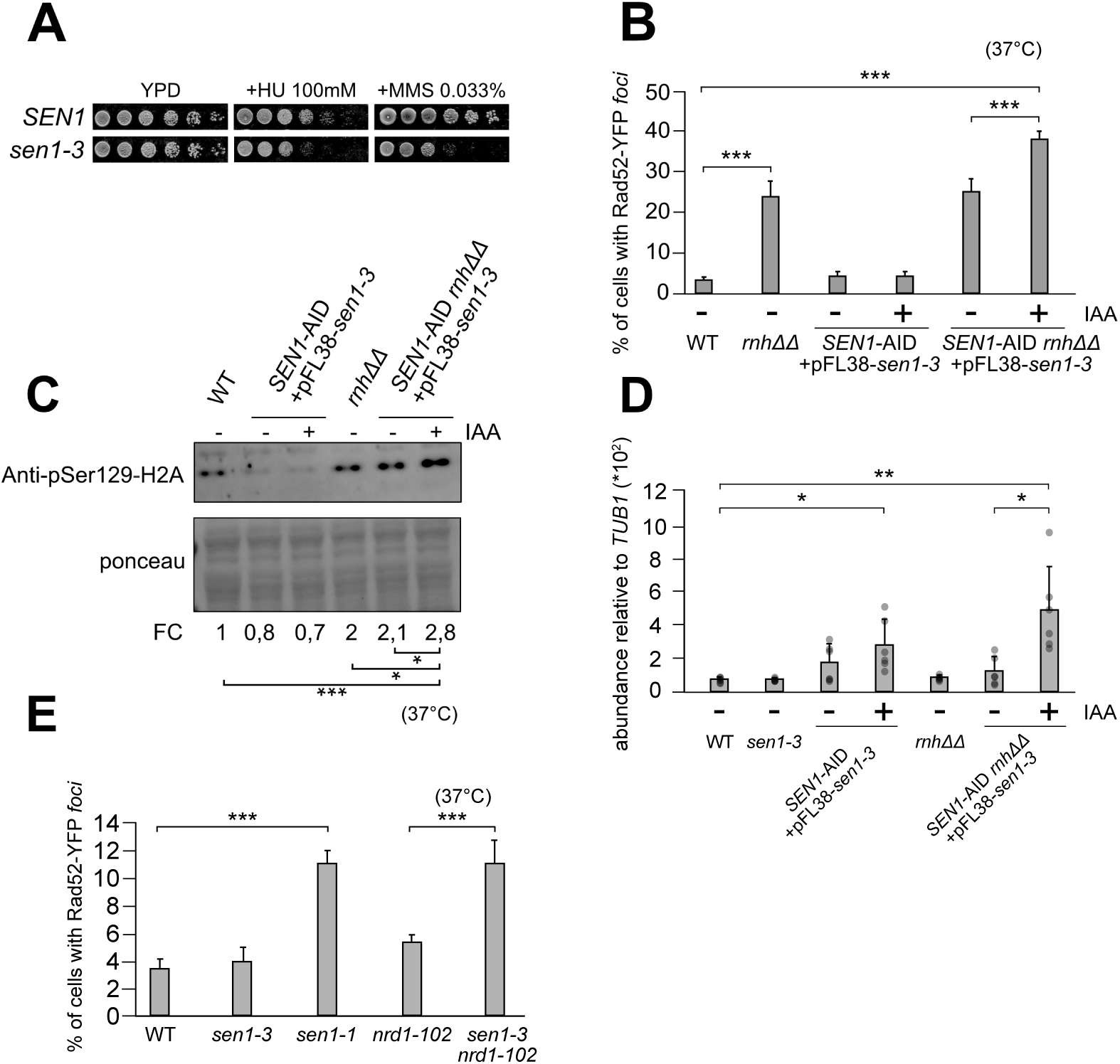
Sen1 cooperates with RNases H to protect genome stability. **A)** Growth assay of *sen1-3* cells and WT controls in the presence or absence of HU and MMS. Plates were incubated for 3 days at 30°C. **B)** Frequency of Rad52-YFP *foci* in asynchronously growing cultures incubated at 37°C for 1 h for the indicated strains, in presence or absence of auxin (IAA) to deplete WT Sen1-AID. **C)** Western blot analyses of the Ser129 phospho form of H2A for cells grown as in B. The average level quantified from three independent replicates is shown at the bottom. D) rDNA copy number assessment by qPCR for cells grown for 3 days at 30°C, in presence or absence of auxin (IAA) to deplete WT Sen1-AID. **E)** As in B. Coupling of the *sen1-3* allele and of transcription termination defects (i.e., *nrd1-102*) leads to levels of DNA damage comparable to the ones observed in *sen1-1* cells. Error bars in B, D and E represent standard deviations; in all panels *p<0.05, **p<0.01, ***p<0.001.

From these observations, we conclude that Sen1 plays important roles in limiting conflicts between RNAPII and RNAPIII at tRNA genes, a role that is dependent on its interaction with RNAPIII but is independent from ongoing replication.

### Roles of Sen1 and RNases H in limiting RNAPII transcription in the ribosomal DNA

Although the ribosomal *loci* are mainly devoted to the production of rRNA by RNAPI and RNAPIII, RNAPII is also found in this region. Because the 5S rRNA is produced by RNAPIII, we first investigated whether the *sen1-3* mutation would affect RNAPII occupancy around the *RDN5* gene. Akin to other RNAPIII transcription units, we clearly observed replication-independent (i.e., observed also in G_1_-arrested cells) RNAPII accumulation antisense of *RDN5* in *sen1-3* cells (Figure S5C). RNAPII accumulation was also observed in *rnh1Δ rnh201Δ* cells suggesting that RNase H is required at this *locus* for efficient RNAPII release. Consistently, R-loops were detected, both by H-CRAC and DRIP (San Martin-Alonso et al., 2021; Wahba et al., 2016) immediately upstream of the paused polymerase (Figure S5C).

Interestingly, we also noticed the occurrence of transcription upstream and antisense of *RDN5* directed towards the close rARS (*ARS1200*) replication origin (Figure 5C), which was only observed in replicating cells and was also RNases H dependent. R-loops were detected by H-CRAC overlapping the whole region of transcription separating the replication origin from the *RDN5* gene, most likely landmarking sites of head-on transcription-replication conflicts. Thus, around *RDN5* Sen1 has replication-dependent and -independent roles in limiting conflicts involving RNAPII transcription.

These findings prompted a closer examination of RNAPII transcription in the rDNA repeats in *sen1-3* and *rnh1Δ rnh201Δ* mutants. Interestingly, we also found in both mutants a region of increased RNAPII occupancy antisense of *RDN37* transcription, roughly corresponding to the internal transcribed spacer 1 (*ITS1*, Figure 5D). This increase was not dependent on replication as it was observed in G_1_-arrested cells and was associated to the formation of R-loops detected with H-CRAC for both Rnh1 and Rnh201 (see Discussion).

We conclude from these data that Sen1 and RNases H play important roles in resolving conflicts involving RNAPII transcription in the ribosomal DNA region.

### RNase H activity and the dual roles of Sen1 transcription termination and in conflict-solving are required for maintaining genome stability

We set up to explore the consequences of the *sen1-3* mutation on genomic stability. We first assessed the sensitivity of *sen1-3* cells to replication stress induced by hydroxyurea (HU) and methyl methanesulfonate (MMS). We found a moderate and strong sensitivity to HU and MMS respectively (Figure 6A), indicating that association of Sen1 with the replisome is required for optimal replication stress response. However, and as we had previously described (Appanah et al., 2020), we did not observe the same genomic instability phenotypes reported for the loss-of-function *sen1-1* mutant, both at the level of Transcription Associated Recombination (Mischo et al., 2011) (Figure S6A), and Rad52 *foci* formation, a hallmark of DSB accumulation and repair (see below, Figure 6E). Also, we did not observe an increase in R-loops by H-CRAC (Figures S4E, S4F and S4G), coherent with previous immunofluorescence analysis of chromosome spreads (Appanah et al., 2020). These differences could be explained if the alterations in the transcription landscape observed in Sen1 loss-of-function but not in *sen1-3* cells (Figure 1) contribute significantly to the genomic instability phenotypes. Also, the functional cooperation with RNases H, underlying the strong genetic interaction with *sen1-3,* might mask phenotypes linked to the loss of interaction with the replisome and RNAPIII. Therefore, we devised a genetic system for analysing the damage induced by the combination of the *sen1-3* and *rnh1Δ rnh201Δ* mutations. We constructed an inducible triple mutant whereby a chromosomal wild-type, AID-tagged Sen1 complements lethality of a *rnh1Δ rnh201Δ* strain containing a plasmid-borne, untagged *sen1-3* allele. Upon addition of auxin (IAA) the wild-type copy is rapidly degraded leaving only the Sen1-3 protein (Mendoza-Ochoa et al., 2019). The induced triple mutant only partially recapitulated the lethality of the *bona fide* triple mutant, possibly because wild-type Sen1 was not fully depleted (Figure S6B), the induced triple mutant phenotype being stronger at 37°C. We monitored the extent of DNA damage by measuring the frequency of Rad52-YFP *foci* (Figures 6B, S6C and S6D). In the non-induced triple mutant, we observed, as expected, the same Rad52 *foci* frequency observed in a double *rnh1Δ rnh201Δ* deletion. Importantly, partial induction of the triple mutant phenotype led to a significant increase in the number of Rad52-YFP *foci* compared to the double RNases H deletion, implying that RNases H and the association of Sen1 with the replisome contribute to limit DNA damage.

Another hallmark of DNA damage, H2A histone phosphorylation at position S129, was monitored by western blot detection. Consistent with the increased Rad52-YFP *foci*, phosphorylation of H2A was found to be significantly increased when the triple mutant, partial phenotype was transiently induced by the addition of IAA (Figures 6C and S6E).

Our genome-wide mapping of transcription-dependent conflicts revealed that Sen1 and RNases H play overlapping functions in limiting conflicts involving RNAPII at the rDNA *locus*. We considered that the DNA damage observed in the triple mutant could at least partially localise to the rDNA. To address this possibility, we analysed the status of the ribosomal array, which experiences changes in the number of repeats upon DNA damage (Kobayashi, 2011). Upon treatment with IAA, all the tested clones of the inducible triple mutant underwent rapid expansion of rDNA copies, suggesting that extensive and generalised damage occurs in the rDNA cluster (Figure 6D). Whether DNA damage is generalized and not only restricted to the rDNA locus is matter for future studies.

Finally, to assess whether the transcription termination defects in Sen1 loss-of-function mutants significantly aggravate the phenotypes linked to the absence of Sen1 at the replisome, we induced a non-coding transcription termination defect in *sen1-*3 cells independently of Sen1 by additionally mutating Nrd1, another component of the NNS complex. We reasoned that if the Sen1 loss-of-function phenotypes were due to a combined termination and conflict-solving defect, a double *nrd1-102 sen1-3* mutant should recapitulate the phenotypes of *sen1-1* cells. This expectation was fully met when analysing the cellular frequency of Rad52 *foci* in the single and double mutants. *Foci* were poorly detected in single mutants, consistently with previous reports (Appanah et al., 2020; Costantino and Koshland, 2018; Mischo et al., 2011), but were found to similar levels in *sen1-1* and *sen1-3 nrd1-102* cells (Figure 6E).

From these results we conclude that by interacting with the replisome and RNAPIII, Sen1 plays important roles at sites of transcription-transcription and transcription-replication conflicts. In the rDNA this role is redundantly exerted by RNases H. In the absence of both enzymes or in the presence of transcription termination defects, extensive DNA damage occurs, which is likely underlying lethality.

## DISCUSSION

Dealing with the extensive occupancy of the genome by transcription events that largely overcome the limits of functional gene annotations is a major challenge in the coordination of concurrent DNA-based activities. In this study we employed high resolution and innovative genomic tools to elucidate the functions of the Sen1 helicase and RNases H in “genomic distancing”. Importantly, this study was performed without altering the physiological transcriptional landscape and in the absence of global replication stress. We propose a model that implicates these enzymes in many sites to control replication- and transcription-transcription conflicts. Importantly, we revisit the role of Sen1 in genomic stability and transcriptional homeostasis by disentangling these functions, whose synthetic association likely impinges on the many phenotypes previously described.

### Loss of Sen1 function generates termination and conflict-solving defects that conjunctly lead to genomic instability

Many earlier studies, including from our laboratory, have clearly established a role for Sen1 in transcription termination of several thousands of non-coding RNA genes (for a review see Porrua et al., 2016). Genome-wide analyses of the effects of Sen1 depletion (this study; Schaughency et al., 2014) demonstrate the induction of major alterations in the non-coding transcriptional landscape, with a large potential for increased interference with concurrent processes. We also describe changes in the gene expression program, which, at least to some extent, parallel the ribosome assembly stress described recently by the Shore, Alberts and Churchman laboratories (Tye et al., 2019; Zencir et al., 2020). These phenotypes are possibly triggered by defective production of snoRNAs, which are major NNS targets and contribute to rRNA maturation, or by defects in transcription termination of rRNA genes, a process in which Sen1 has been implicated (Kawauchi et al., 2008). Together, these results demonstrate that the transcription and gene expression program of Sen1 loss-of-function mutants is largely distinct from the one of wild-type cells, and raise questions about the assessment of genomic instability phenotypes in such non-physiological conditions.

It has been shown that alterations in transcription termination generated with other NNS mutants (i.e., *nrd1* and *nab3*) do not induce instability *per se* (Alzu et al., 2012; Costantino and Koshland, 2018; Mischo et al., 2011). These earlier findings indicate that transcription termination defects are not sufficient, alone, to generate genomic instability. However, it is possible that when termination alone is impaired (e.g. in *nab3 or nrd1* mutants), the increased transcriptional challenge at sites of conflicts can still be resolved by Sen1 and RNases H, while in the additional absence of this conflict-resolving function increased DNA damage occurs or is not repaired efficiently. Fully consistent with this notion is our finding that impairing transcription termination independently of Sen1 (i.e., by mutation of Nrd1) in *sen1-3* cells, fully recapitulates the DNA damage phenotype of *sen1-1* cells, which are defective for both termination and conflict-solving functions of Sen1 (Figure 6).

### On the physiological relevance of Sen1 binding to the replisome

Focal to our study is the *sen1-3* mutant, which we have previously shown to fully lose interaction with replisome components (Appanah et al., 2020) and with RNAPIII (Xie et al., 2021). Our working hypothesis underlying this study has been that the interaction of Sen1 with the replisome allows either its recruitment or its function at sites where transcription collides with replication to dismantle the elongation complex for giving way to replication.

Because *sen1-3* cells have no major growth or hyper-recombination phenotypes (Figures 6 and S6), it would be legitimate to question the physiological relevance of the functions impaired in this mutant. However, the many genetic and biochemical observations reported in our study support the notion that the *sen1-3* mutation affects important facets of Sen1 function for which redundant mechanisms are in place that mask the effects of the single mutation.

In this perspective, while this work was in progress it was reported that DNA damage and fork stalling occur under Sen1 depletion at sites of TRCs specifically under conditions of HU-induced replication stress (Zardoni et al., 2021). In this study, the expression of genes that host TRCs was also found to be altered, a phenotype that we did not observe even upon Sen1 depletion (data not shown). Our interpretation for these differences is that failure to complete transcription at some genes hosting TRC sites results from a synergistic effect of decreased replication fork progression due to the HU treatment and the failure to efficiently remove RNAPII by Sen1 at TRCs, which together delays the resolution of conflicts and affect the resumption of novel transcription cycles.

We provide evidence that Sen1, by binding to the replisome, releases RNAPII engaged in conflicts with replication in the very 5’-end of genes and in other regions of pausing, suggesting that it is RNAPII pausing and not the 5’-end of genes *per se,* that determines the preferential sites of conflicts. Our data suggest that RNases H do not cooperate with Sen1 in the TSS-proximal locations, most likely because the limited window of free nascent RNA, which is also occupied by the capping complex, does not allow efficient formation of R-loops. A similar, genic RNAPII accumulation was described in *dicer* mutants in the 3’-end of *S. pombe* genes, which was also proposed to be due to the defective resolution of TRCs (Castel et al., 2014). The reasons for the difference in the position of RNAPII accumulation are not clear, but they might be due to a different mechanism of action of Sen1 and Dicer or to the different distribution of RNAPII pausing at *S. pombe* genes.

While this work was in preparation, a similar increase in RNAPII occupancy was reported to occur in cells expressing loss-of-function mutants of the human Sen1 homologue *SETX* or in *ΔSETX* cells (Kanagaraj et al., 2022). Although it was not shown whether these phenotypes are due to TRCs, they were found more prominently in very long (>100kb) genes, which generally host common fragile sites (CFS). Genes hosting CFS were also shown in this study to be frequently subject to genomic rearrangements in the absence of SETX. Thus, the role of Sen1 in solving TRCs within genes might be conserved for human SETX, and could also be independent from its function in terminating non-coding transcription, which is not clearly established in human cells.

### On the relationship between Sen1 and R-loops

A salient question, central to this study, concerns the functional relationships between Sen1 and RNases H. Although both classes of enzymes have been involved in the resolution of R-loops and have been proposed to work together at these structures (Costantino and Koshland, 2018), we could not globally detect increased R-loop levels in *sen1-3* cells by H-CRAC (Figures S4E-G), chromosome spreads (Appanah et al., 2020), or by DRIP-qPCR at a few R-loop-prone sites (data not shown). This indicates that the interaction of Sen1 with the replisome is not required for suppressing R-loops and might suggest that the hybrids increase detected in *sen1* lack-of-function cells is also linked to the defective transcriptional termination phenotype. Because R-loops are known to be sites prone to DNA damage, this notion is consistent with the finding reported here that increased sites of DNA damage are observed when associating a termination defect generated by the *nrd1-102* allele to the *sen1-3* mutation. In this perspective, it is possible that Sen1, rather than unwinding R-loops that are constitutively formed, prevents their formation by a dual action in restricting non-coding transcription and solving conflicts with replication. In agreement with this hypothesis, we have previously reported biochemical and single-molecule evidence that purified Sen1 is poorly processive in unwinding DNA:RNA duplexes (Han et al., 2017; Wang et al., 2019).

### Sen1 and RNases H cooperate to release RNAPII at the ribosomal DNA

At the rRFB and upstream of *RDN5,* both the *sen1-3* and the *rnh1Δ rnh201Δ* mutations lead to replication-dependent, increased RNAPII occupancy, suggesting that Sen1 and RNases H play complementary or redundant functions in preventing RNAPII transcription from invading the rDNA, which could cause genomic instability in the region. Indeed, it was previously shown that forcing transcription from the E-pro region using the strong *GAL10* promoter releases cohesins and induces rDNA amplification in a strain containing only two rDNA repeats (Kobayashi and Ganley, 2005). Consistent with this notion is our finding that induction of the triple *sen1-3 rnh1Δ rnh201Δ* is associated to an increased level of rDNA repeats (Figure 6D). It is also possible that efficiently removing RNAPII from upstream of the stalled fork is important to limit replication stress and favor the progression of replication in this region where two replication forks converge.

Our findings might suggest a role of RNases H in transcription termination. Mechanistically (Figure 7), the removal of R-loops by RNases H might weaken the stability of the elongation complex, for instance if, upon pausing, RNAPII would backtrack on a substrate on which RNase H has previously removed the DNA-associated nascent transcript engaged in an R-loop. Digestion of the DNA:RNA heteroduplex by RNase H might also facilitate the access of Sen1 to the nascent RNA close to the stalled elongation complex and induce termination. This would be consistent with the notion that DNA:RNA or RNA:RNA double stranded regions of the nascent transcript affect Sen1-dependent termination both *in vitro* (Porrua and Libri, 2013) and *in vivo* (Xie et al., 2021), possibly by affecting binding or translocation on the nascent transcript (Porrua and Libri, 2013; Wang et al., 2019).

**Figure 7:**
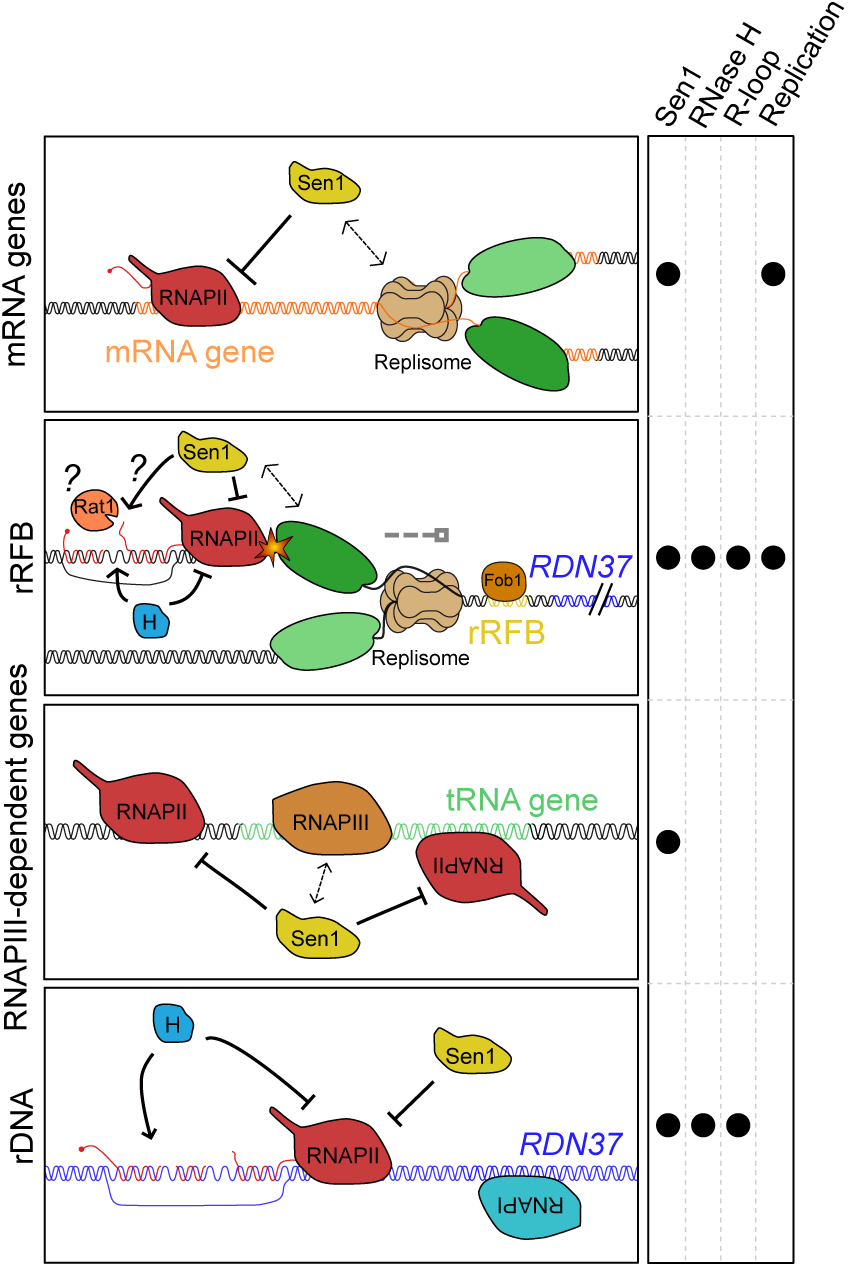
Model of the function of Sen1 and RNases H in limiting RNAPII-driven conflicts. Sen1 resolves transcription-replication and transcription-transcription conflicts by dislodging RNAPII from several locations including the 5’-end of coding genes, the rRFB, the rDNA and tRNA genes. During TRCs Sen1 is recruited via interaction with the replisome, while at tRNA genes the recruitment is achieved via interaction with RNAPIII. RNases H assist Sen1 in limiting RNAPII at the rRFB and in the rDNA, via a mechanism that could involve Rat1.

One interesting alternative is that degradation of the R-loop-engaged RNA might generate an entry site for the Rat1 exonuclease, which would degrade the 3’ portion of the nascent RNA and terminate transcription by the “torpedo” mechanism (Porrua et al., 2016). Consistent with this notion, we found that a double *rat1-1 sen1-3* mutant has a strong synthetic growth defect (Figure S4H, compare growth of *rat1-1* to *rat1-1 sen1-3* cells) that phenocopies the genetic interaction of *sen1-3* with the double *rnh1Δ rnh201Δ,* possibly suggesting that Rat1 is a downstream effector of RNases H. This would also be in agreement with a recent report showing that cleavage of the nascent RNA by oligonucleotide-directed digestion (possibly mimicking R-loop digestion) can induce transcription termination (Lai et al., 2020).

### Sen1 resolves transcription-transcription conflicts

We show that Sen1 solves conflicts between RNAPII and RNAPIII (Figures 5 and S5), by virtue of its interactions with the latter that is also lost in *sen1-3*, and, consistently, RNAPII accumulation is observed also in the absence of replication. The fact that the same region of Sen1 mediates alternative interactions with RNAPIII and the replisome might have important functional implications, to ensure that the two functions of the helicase, although mechanistically similar, remain distinct, and that replisome components and RNAPIII are never found, inappropriately, in the same complex, connected by Sen1. Perhaps this region of Sen1 mediates contacts with other molecular machineries to exert similar functions. In this regard, we observed marked RNAPII persistence in *sen1-3* cells antisense of the *RDN37* transcription unit, in correspondence of the *ITS1*. Limiting RNAPII in this region is also dependent on RNases H and prominent levels of R-loops are observed by H-CRAC. However, this accumulation was also observed in the absence of replication, and why it occurs in the *sen1-3* mutant is presently unclear. One possibility is that the interaction with another factor, responsible for recruiting Sen1 at this site, is also lost in the *sen1-3* mutant and that RNAPII persists at sites of head-on conflicts with RNAPI. It is enticing to speculate that Sen1 is recruited to the nucleolar site of rRNA transcription by RNAPIII, while transcribing the 5S rRNA or tRNAs that are also transcribed in clustered nucleolar regions (Thompson et al., 2003; Haeusler and Engelke, 2006). Loss of Sen1 from RNAPIII complexes in *sen1-3* cells would also bring about defective management of RNAPI-RNAPII conflicts.

In the light of the impact of the *sen1-3* mutation on RNAPIII termination, we considered the possibility that the accumulation of RNAPII at the rRFB might be linked to its role at the upstream *RDN5* gene. However, RNAPII accumulation occurs in a position that is clearly downstream of the region of RNAPIII readthrough, as shown by RNAPIII CRAC analyses (Figure S4A).

Together, these results, obtained in the absence of possibly interfering transcriptional defects, allow attributing to Sen1 a role of “key regulator of conflicts” that is similar in many aspects to the one described for Dicer in *S. pombe* (Castel et al., 2014). Although the mechanism of action of the two factors is unlikely to be similar, converging evolution might have hijacked existing cellular mechanism to fulfill the important role of coordinating essential cellular processes.

### H-CRAC is a suitable method for R-loops detection genome-wide

We describe here a novel method to detect R-loops with unprecedented sensitivity and resolution. H-CRAC meets all the essential landmark requirements we assessed for reliable R-loop detection (Chédin et al., 2021) strongly supporting the notion that RNase H targets detected by this method represent *bona fide* R-loops. It is important to stress that H-CRAC is fundamentally different from ChIP as it detects the interaction of RNases H with the RNA and is not expected to sense recruitment of the enzyme to the DNA in the absence of a specific contact with its targets.

Our maps are similar to published DRIP-seq maps (Figure 4B). Nevertheless, in many cases differences are also observed, which might be due to the better resolution and most likely higher sensitivity of H-CRAC relative to DRIP-seq (Figure 4A), but also to differences in the detection potential of the two methods. By possibly detecting inherently different targets, the two methods might provide overlapping and complementary outputs.

The genome-wide analyses of R-loop distribution in yeast by H-CRAC will be presented in a separate report, but to the light of the results and controls presented here we trust that H-CRAC will provide invaluable information to study R-loop biology and the relationships with transcription and genome maintenance.

## ACKNOWLEDGEMENTS

We wish to thank Andres Aguilera, Doug Koshland, Rodney Rothstein for providing strains or plasmids; Giacomo De Piccoli, Sarah Lambert, Vincent Vanoosthuyse, Michel Werner, Frédéric Chédin, Julien Gros and the members of the Libri and Palancade labs for critical reading of the manuscript and fruitful discussion; Vasudha Sharma for help with the generation of mutants and with microscopy experiments. We also wish to thank Yan Jaszczyszyn for expert technical help in preparing CRAC libraries for sequencing. This work has benefited from the facilities and expertise of the high throughput sequencing core facility of I2BC (Centre de Recherche de Gif - http://www.i2bc-saclay.fr/).

## FUNDING

This work was supported by the Centre National de la Recherche Scientifique (C.N.R.S.), the Fondation pour la Recherche Medicale (F.R.M., programme Equipes 2019 to D.L.), l’Agence National pour la Recherche (ANR-16-CE12-0022-01 to D.L. and P.P.; ANR-21-CE12-0040-01 to D.L. and B.P.; ANR-18-CE12-0003 to B.P.; ANR-19-CE12-0016-01 to P.P.), the Fondation ARC pour la Recherche sur le Cancer and the Ligue Nationale contre le Cancer to B.P. and the IdEx Université de Paris (ANR-18-IDEX-0001 to B.P.).

U.A. was supported by the French Ministry for Education and Research, by the Fondation ARC pour la Recherche sur le Cancer and the EUR G.E.N.E. (reference #ANR-17-EURE-0013), which is part of the Université de Paris IdEx #ANR-18-IDEX-0001 funded by the French Government through its “Investments for the Future” program.

## AUTHOR CONTRIBUTIONS

U.A. and D.L. designed experiments. U.A. performed all the experiments and analysis excluding the raw data analysis of the DNA Copy number and the Sen1-AID RNAPII CRAC and the ensuing bioinformatic analysis presented in Figure 1. D.C. performed the Sen1-AID RNAPII CRAC. G.W. offered technical assistance for several experiments. A.L. assisted U.A. with the DNA Copy Number analysis. B.P. assisted U.A. with DRIP experiments and Rad52 *foci* imaging. RA provided the sen1-3 strain ahead of publication. D.L. performed the bioinformatic analysis presented in Figure 1. U.A. and D.L. wrote the manuscript. D.L. supervised the project. D.L., B.P. and P.P. collected funding. All authors provided feedbacks during the writing and approved the manuscript.

## DECLARATION OF INTERESTS

The authors declare no competing interests.

## MATERIALS AND METHODS

## RESOURCE AVAILABILITY

### Lead contact

Further information and requests for resources and reagents should be directed to and will be fulfilled by the lead contact, Domenico Libri.

### Materials availability

This study did not generate new unique reagents.

### Data and code availability

All the datasets generate by this study have been deposited in the Geo Expression Omnibus repository at NCBI and are available using the code GSE195936. Any additional information required to reanalyze the data reported in this paper is available from the lead contact upon request.

## EXPERIMENTAL MODEL AND SUBJECT DETAILS

*Saccharomyces cerevisiae* is the model organism used in this study. All yeast strains used in this study are listed in Key Resource Table. All strains derive from W303 or BMA64, which is a *trp1Δ* derivative of W303. Strains newly modified were constructed with standard procedures (Longtine et al., 1998).

## METHOD DETAILS

### Transcription-Associated Recombination (TAR) assay

To assess the frequency of recombination, strains of interest were transformed with the pRS314-L recombination reporter at 30°C. Recombination events were scored by assessing the number of cells containing a functional, recombined *LEU2* gene relative to the total number of cells plated. Six colonies of at least three independent transformants were analysed.

### Cell growth for CRAC and Copy Number experiments

For each condition, 2 L of cells expressing an HTP-tagged version of the protein of interest expressed either from the endogenous locus (i.e., Rpb1, Rnh1, Rnh201) or from a plasmid (hsRNH1) were grown in logarithmic phase to OD_600_=0.6 at 30°C in a CSM-TRP medium. Cells ectopically expressing the mAIRN construct were grown in CSM-Trp-Ura. Cells over-expressing hsRNH1 were grown in CSM-Trp-His.

In conditional depletion experiments, Indole-3’-Acetic Acid (IAA, Sigma) was supplemented at a final concentration of 500 μM 1 h before UV treatment.

G_1_ cell cycle arrest was triggered at OD_600_=0.3 by 3 consecutive additions of 4, 8 and 4 mg of *α*-factor spaced by 40 min. UV treatment was performed 40 min after the last addition of *α*-factor.

For analyses in S-phase, cells were arrested in G_1_ by *α*-factor as described above and released into S-phase by removing *α*-factor by filtration on a glass microfiber filter (pore ⌀=1.6 μm). Cells were washed while still on the filter and then resuspended in 2 L of fresh medium lacking *α*-factor at 30°C for 30 min or 45 min before UV treatment.

G_1_ arrest and S-phase release were systematically verified by visualisation of cell morphology and flow cytometry (Figure S2J). Two biological replicates were performed for each condition, showing high correlation (Fig S7B).

### Assessment of rDNA copies by quantitative PCR (qPCR)

Independent clones of the indicated strains were grown for 3 days on a YPD plate at 30°C before being used for DNA isolation by standard phenol:chroloform extraction, followed by RNase treatment. DNA amounts were quantified by real-time PCR with a LightCycler 480 system (Roche) using SYBR Green incorporation according to the manufacturer’s instructions. Oligos DL1561 and DL1562 were used to amplify a region in the rDNA *locus* while oligos DL377 and DL378 were used to amplify a region in *ACT1* used as a reference for a unique DNA sequence in the genome.

### Chromatin Immunoprecipitation (ChIP)

Exponentially growing cells were crosslinked with formaldehyde in the presence of 1% formaldehyde for 10 min at 30°C. For conditional depletion of Fob1, Indole-3’-Acetic Acid (IAA, Sigma) was supplemented at a final concentration of 500 μM 2 h before crosslinking. After quenching with 100 mM glycine, the cell pellets were lysed by beads beating in 1 ml of lysis buffer (50 mM Hepes pH 7.5, 140 mM NaCl, 1 mM EDTA, 1% triton X-100, 0.1% deoxycholate, 1X protease inhibitors cocktail, complete EDTA-free, Roche). Cell lysates were sonicated using a Bioruptor (Diagenode). A volume of extract roughly corresponding to 2 OD_600_ was supplemented with 0.5% BSA and 50 μg/mL salmon sperm DNA and immunoprecipitated using affinity-purified with M-280 tosylactivated dynabeads coupled with rabbit IgGs for 1 h at 4°C. Washes were as follow: twice with lysis buffer, twice with lysis buffer containing 360 mM NaCl; twice with 10 mM Tris pH 8, 250 mM LiCl, 0.5% Nonidet-P40, 0.5% deoxycholate, 1 mM EDTA and once with 10 mM Tris–HCl pH 8, 1 mM EDTA. Elution was performed in the presence of 50 mM Tris pH 8, 10 mM EDTA, 1% SDS for 10 min at 65°C. The eluates were deproteinized with proteinase K (Sigma, 0.2 mg/mL) and uncrosslinked for 30 min at 65°C. Immunoprecipitated DNAs were purified with the Qiaquick PCR purification kit (Qiagen). Input and eluate DNAs were quantified by real-time PCR with a LightCycler 480 system (Roche) according to the manufacturer’s instructions. Oligos DL4942 and DL4943 were used to amplify the rDNA region upstream Fob1 biding site, and precisely where RNAPII accumulation occurs as detected by CRAC.

### UV-Crosslinking and cDNA analysis (CRAC)

The CRAC protocol used in this study is derived from Granneman et al. (2009) with some modifications described in (Challal et al., 2018; Colin et al., 2014).

Briefly, cells were crosslinked by UV exposure for 50 seconds using a W5 UV crosslinking unit (UVO3 Ltd) and harvested by centrifugation at 4°C. Cell pellets were washed once with ice-cold 1x PBS, weighted and resuspended in 2.4 mL/(g of cells) of TN150 buffer (50 mM Tris pH 7.8, 150 mM NaCl, 0.1% NP-40 and 5 mM *β*-mercaptoethanol) supplemented with fresh protease inhibitors (AEBSF, Complete™ EDTA-free Protease Inhibitor Cocktail, Roche). Emulsions were snap-frozen in droplets in liquid nitrogen and cells subjected to cryogenic grinding using a Ball Mill MM 400 (5 cycles of 3 minutes at 20 Hz). The resulting frozen lysates were thawed on ice, treated with DNase I (165 units per gram of cells) incubated at 25°C for 1h to solubilize the chromatin and then clarified by centrifugation at 16 krpm for 30 min at 4°C.

RNA-protein complexes were affinity-purified with M-280 tosylactivated dynabeads coupled with rabbit IgGs (10 mg of beads per sample), washed with TN1000 buffer (50 mM Tris pH 7.8, 1 M NaCl, 0.1% NP-40 and 5 mM β-mercaptoethanol), and eluted by TEV protease digestion. RNAs were subjected to partial degradation by treating with 0.2 U of Rnase cocktail (Rnace-IT, Agilent) and the reaction was stopped by the addition of guanidine–HCl to a final concentration of 6 M. Eluates underwent then a second immobilisation on Ni-NTA columns (Qiagen, 100 μl of slurry per sample) overnight at 4°C and were extensively washed. Sequencing adaptors were ligated to the RNA molecules as described in the original procedure. RNA-protein complexes were eluted with elution buffer containing 50 mM Tris pH 7.8, 50 mM NaCl, 150 mM imidazole, 0.1% NP-40 and 5 mM β-mercaptoethanol fractionated using a Gel Elution Liquid Fraction Entrapment Electrophoresis (GelFree) system (Expedeon) following manufacturer’s specifications. The fractions containing the protein of interest were treated with 100 μg of proteinase K, and RNAs were purified and reverse-transcribed using reverse transcriptase Superscript IV (Invitrogen).

After quantification of the recovered material via quantitative PCR, the cDNAs were amplified with an appropriate number of PCR cycles using LA Taq polymerase (Takara), and then the reactions were treated with 200 U/mL of Exonuclease I (NEB) for 1 h at 37°C. Finally, the DNA was purified using NucleoSpin columns (Macherey-Nagel) and sequenced on a NextSeq 500 Illumina sequencer.

The H-CRAC protocol contained a few modifications to improve the recovery of tagged RNases H. DNase I treatment was replaced by a step of chromatin shredding by sonication in an ice-cold bath (15 min, High, 45 sec ON/OFF, Diagenode). The GelFree fractionation was omitted to avoid loss of material because the eluate after the second purification step was judged sufficiently pure.

### Copy Number analysis

An aliquot roughly corresponding to 0.2 g of lysate powder from the CRAC experiment was transferred to a separate tube and used to perform genomic extraction using the Genomic-tip 20/G kit (QIAGEN) following manufacturer’s specifications. DNA was fragmented using sonication (∼200 to 500 bp size range). Sequencing libraries were prepared using a ThruPLEX DNA-seq kit (Rubicon Genomics). Next-generation sequencing was performed on a HiSeq 4000 (Illumina). Single-end reads of 50 bp were aligned to the *S. cerevisiae* genome (2011).

### DNA:RNA immunoprecipitation (DRIP)

DNA:RNA hybrid immunoprecipitation (DRIP) was performed using the S9.6 DNA:RNA hybrid-specific monoclonal antibody according to a published procedure (Mischo et al., 2011; Wahba et al., 2016), with the modifications described in (Bonnet et al., 2017). Briefly, genomic DNA was phenol-extracted from cells growing exponentially and isolated by ethanol precipitation. 50 µg of purified nucleic acids were digested by a cocktail of restriction enzymes (EcoRI, HindIII, XbaI, SspI, BsrGI; FastDigest enzymes; Thermo Scientific) for 30min at 37°C in a total volume of 100 µL. An RNase H treatment (10 units, Sigma) was included in the restriction reaction of control samples to assess the specificity of the DRIP signal. Digested samples were further diluted 4-fold with FA1 buffer (0.1% SDS, 1% Triton, 10 mM HEPES pH 7.5, 0.1% sodium deoxycholate, 275 mM NaCl) and incubated overnight at 4°C in the presence of 1.5 µg of S9.6 purified antibody (Kerafast). Antibody-associated DNA:RNA hybrids were then captured on protein G Sepharose beads (GE Healthcare), washed and purified according to standard ChIP procedures. Input and immunoprecipitated DNA amounts were quantified by real-time PCR with a LightCycler 480 system (Roche) using SYBR Green incorporation according to the manufacturer’s instructions. Oligos DL4519 and DL4520 were used to amplify the mAIRN *locus* while oligos DL4597 and DL4598 were used to amplify an intergenic region located in proximity to the *HO* gene and used a negative control. The amount of DNA in the immunoprecipitated fraction was divided by the amount detected in the input to evaluate the percentage of immunoprecipitation (% of IP).

### Imaging of Rad52-YFP *foci*

Rad52-YFP *foci* formation was assessed in exponentially growing cells (0.5 ≤ OD_600_ ≤ 1) in CSM medium at 30°C or 37°C as indicated. For wild-type Sen1-depleted conditions, Indole- 3’-Acetic Acid (IAA, Sigma) was supplemented at a final concentration of 500 μM 1 h before imaging. Wide-field fluorescence images were acquired using a Leica DM6000B microscope with a 100X/1.4 NA (HCX Plan-Apo) oil immersion objective and a CCD camera (CoolSNAP HQ; Photometrics). The acquisition system was piloted by the MetaMorph software (Molecular Devices). For all images, z stacks sections of 0.2 μm were acquired using a piezo-electric motor (LVDT; Physik Instrument) mounted underneath the objective lens. Images were scaled equivalently and z-projected using ImageJ. An average of three experiments, each of them visualizing at least 300 cells per condition, is shown in the figures.

For thermosensitive alleles, and their relative controls, after reaching exponential growth cells were shifted to 37°C by the addition of pre-warmed media, and incubated for 1 h before imaging.

### Protein analyses

Proteins levels were analysed using current methodologies.

## QUANTIFICATION AND STATISTICAL ANALYSIS CRAC

CRAC datasets were analysed as described (Candelli et al., 2018; Challal et al., 2018). The pyCRAC script pyFastqDuplicateRemover was used to collapse PCR duplicates using a 6 nucleotides random tag included in the 3’ adaptor (see Key Resources Table). The resulting sequences were reverse complemented with Fastx reverse complement (part of the fastx toolkit, http://hannonlab.cshl.edu/ fastx_toolkit/) and mapped to the R64 genome (Cherry et al., 2012) with bowtie2 (-N 1) (Langmead and Salzberg, 2012).

Reads aligning to regions corresponding to very abundant mature RNAs such as tRNAs or rRNAs were systematically masked because susceptible of deriving from contamination. Although the RNAPII complex has undergone several purification steps, a significant amount of mature tRNAs and rRNAs remain (to a lower level than in techniques that do not use crosslinking).

For experiments in which overexpression of human RNH1 was used a mean to reduce R-loops level, data were normalised using a *S. pombe* spike-in that was added to the *S. cerevisiae* cultures before UV-crosslink.

### Quantification and statistical analysis

The vast majority of the analyses were performed with inhouse scripts in the R studio environment. Sen1-AID CRAC datasets were analysed using the Galaxy web platform at usegalaxy.org (Afgan et al., 2018).

For all RNAPII CRAC data, the working group of 3000 genes with the highest expression level was selected by computing HT-seq count normalised to the size of the gene. This allowed excluding from our analysis genes with very low or background signal, which are potential source of computational biases.

The skew index was defined as the ratio between the RNAPII CRAC signals in the windows [0; +200]/[+300; +500] relative to the TSS for each gene. A *sen1-3*/WT skew index ratio was used to select genes with increased 5’-end RNAPII accumulation in the mutant relative to the wild-type. All features with a skew index ratio exceeding the mean plus one standard deviation of the distribution were considered to be affected.

For Copy Number analysis, all regions with a score >1 were considered as undergoing replication. For the selection of the “already replicated genes” genes overlapping with a replicated region with a copy number score exceeding the 95^th^ percentile were chosen.

When average values were represented, error bars indicate standard deviation. T tests were used to compare distributions and p-values are indicated.

## RESOURCE TABLE

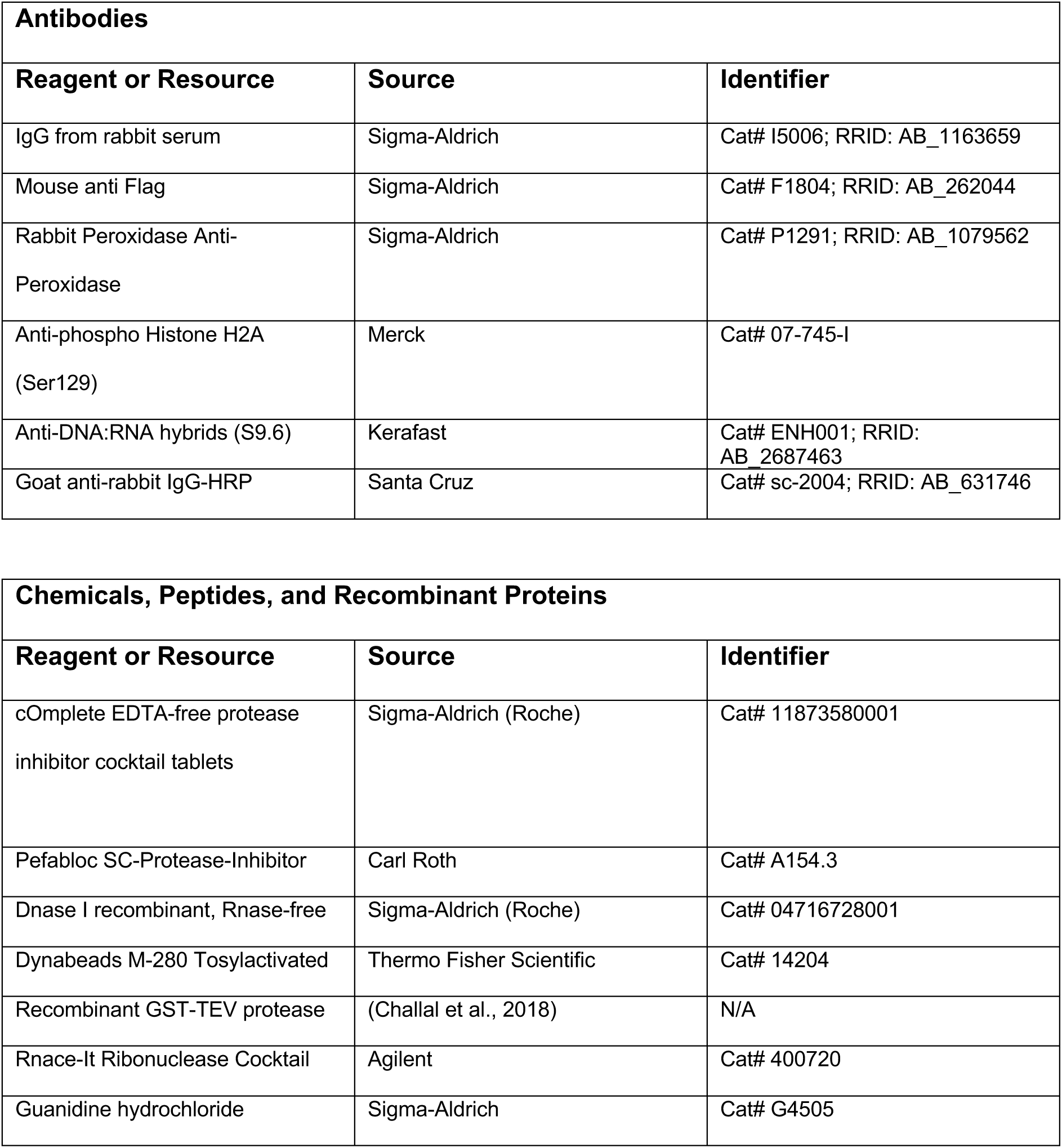

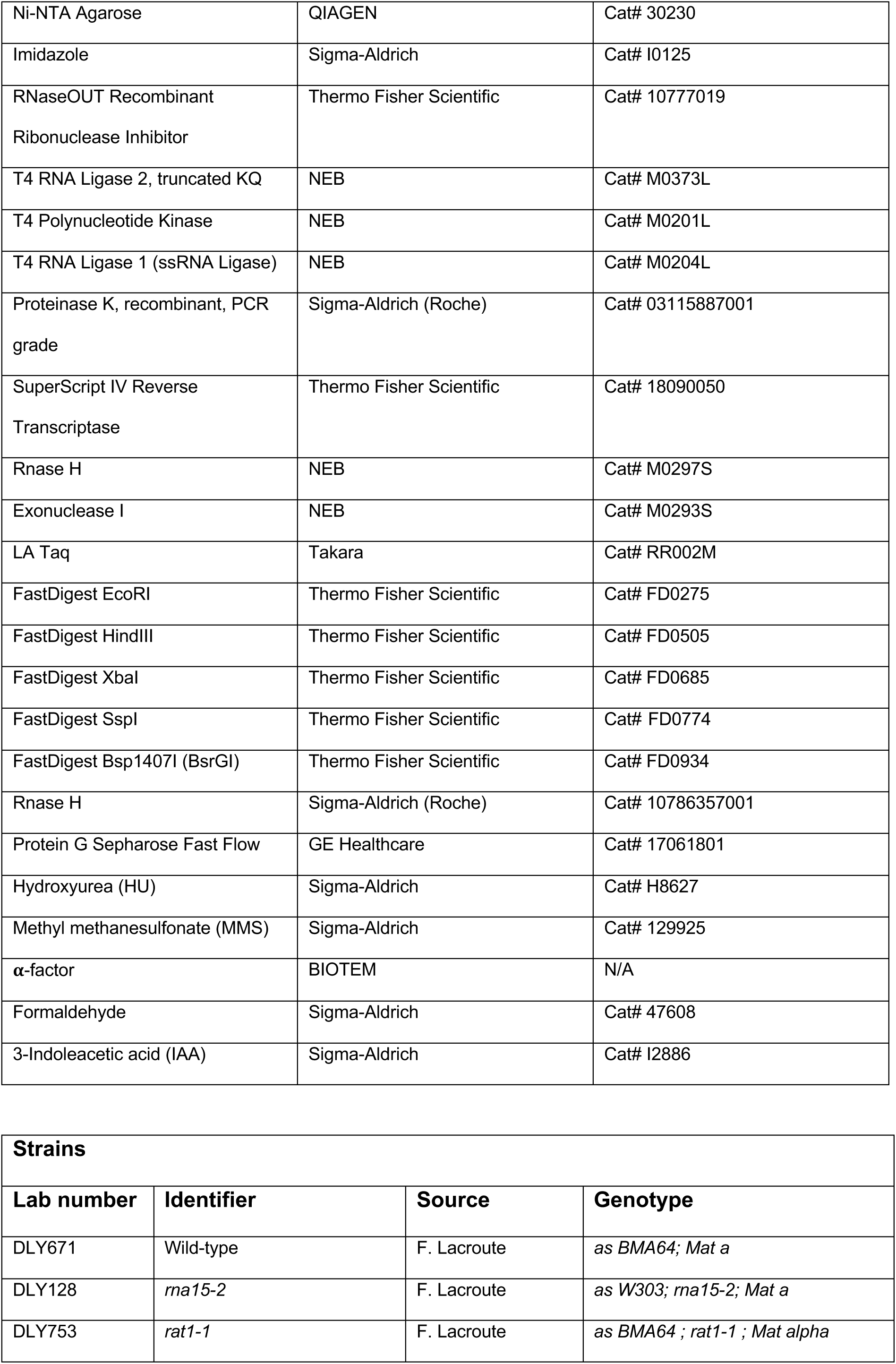

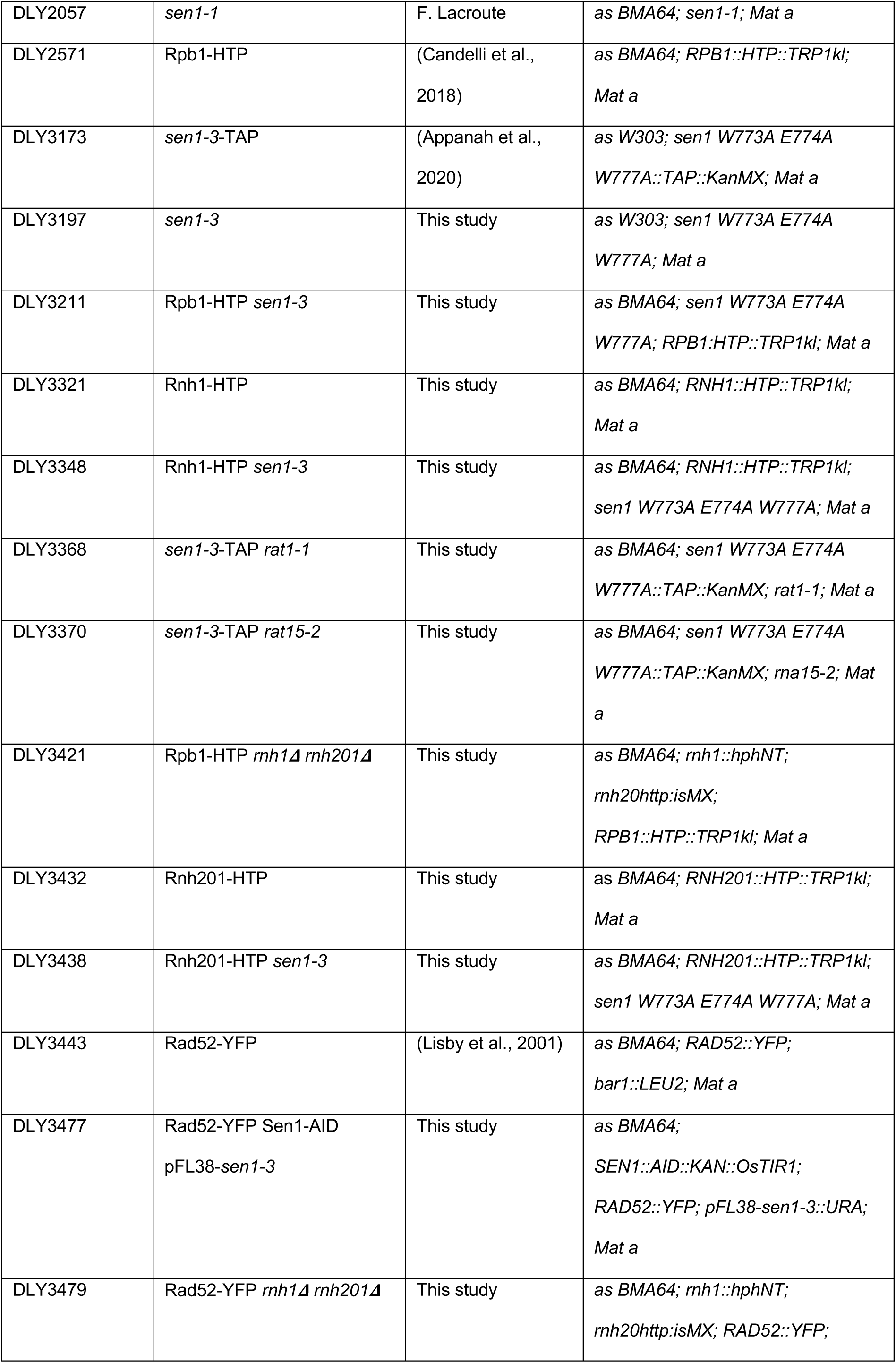

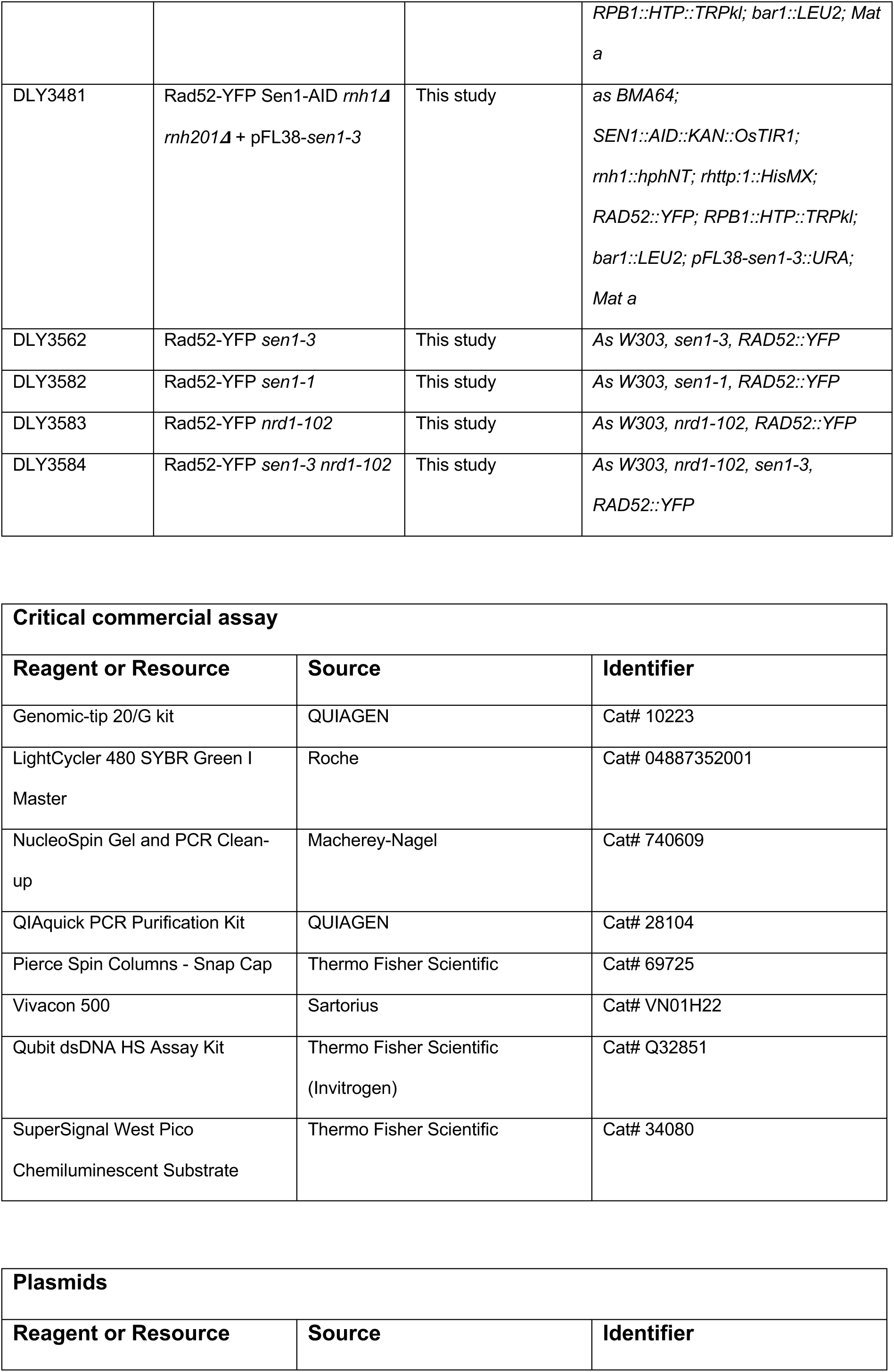

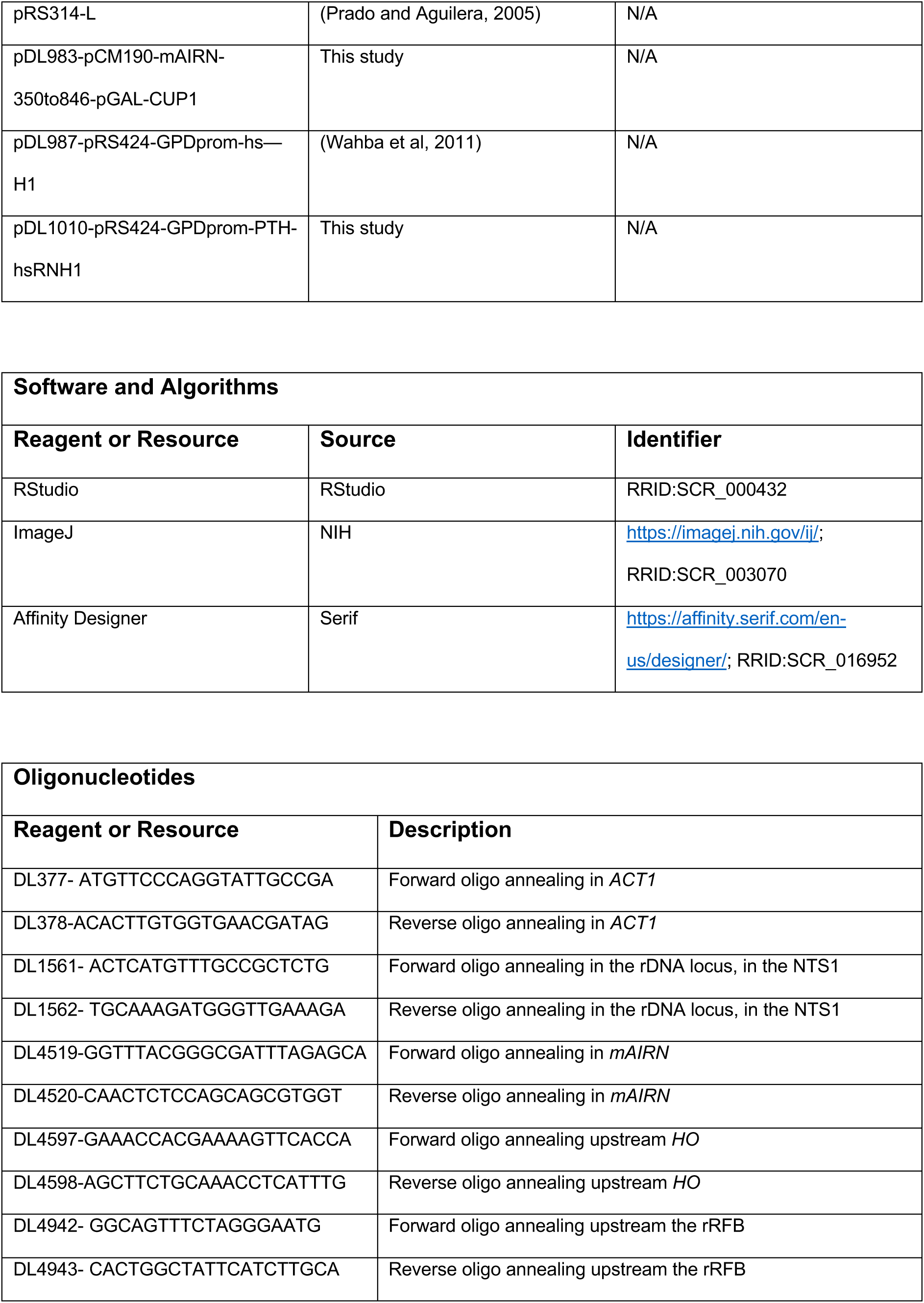

## SUPPLEMENTAL FIGURES

**Supplementary Figure 1 related to Figure 1:**
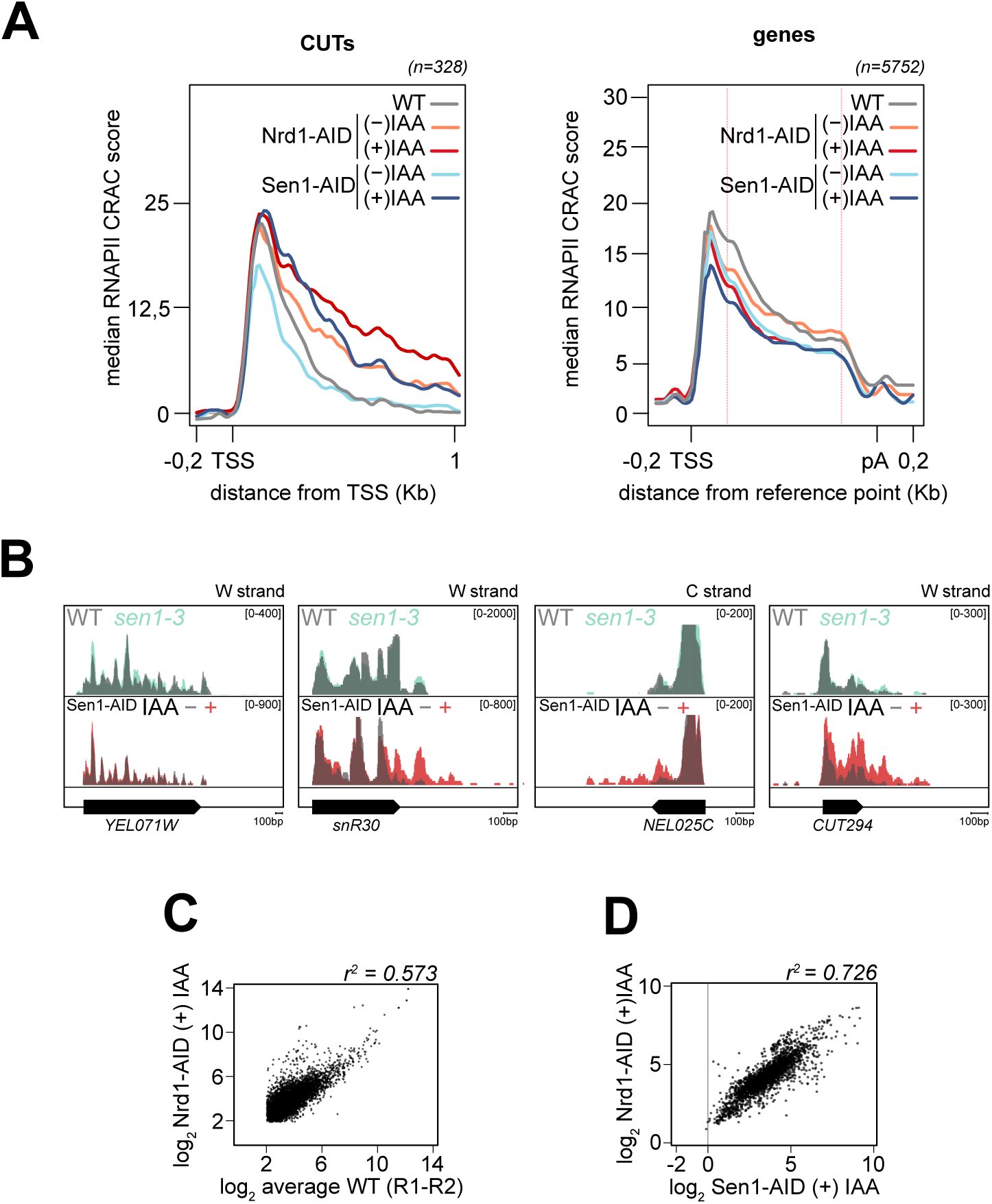
**A)** Metagene analysis of RNAPII distribution at a subset of validated Cryptic Unstable Transcripts (CUTs, left panel) and at mRNA-coding genes (right panel) in wild-type (WT), Nrd1-AID and Sen1-AID cells in presence or absence of auxin (IAA). The CUTs plot is unscaled and aligned at the Transcription Start Site (TSS); The genes plot is aligned on the TSS and pA site, but unscaled in the 400 nt windows centered on each alignment point. Values on the y-axis correspond to the median coverage. Note that the 3’-end of CUTs is not well defined, hence the increase in RNAPII occupancy upon depletion of Nrd1 or Sen1 is spread over a large region. **B)** Representative snapshots illustrating the absence of termination defects at coding and non-coding genes in *sen1-3* cells. For comparison, the tracks derived from Sen1-AID cells in presence or absence of auxin (IAA), showing read-through at CUTs and snoRNAs are also included. *YEL071W* is shown as a representative example of lack of termination defects at coding genes in Sen1-AID cells even in presence of IAA. **C)** Scatter plot as in Figure 1A but for Nrd1-AID strain under depletion conditions (auxin added for 1 hour). **D)** Scatter plot analysis showing the good correlation of RNAPII CRAC density signals in mRNA-coding genes in Sen1- and Nrd1-depleted cells.

**Supplementary Figure 2 related to Figure 2:**
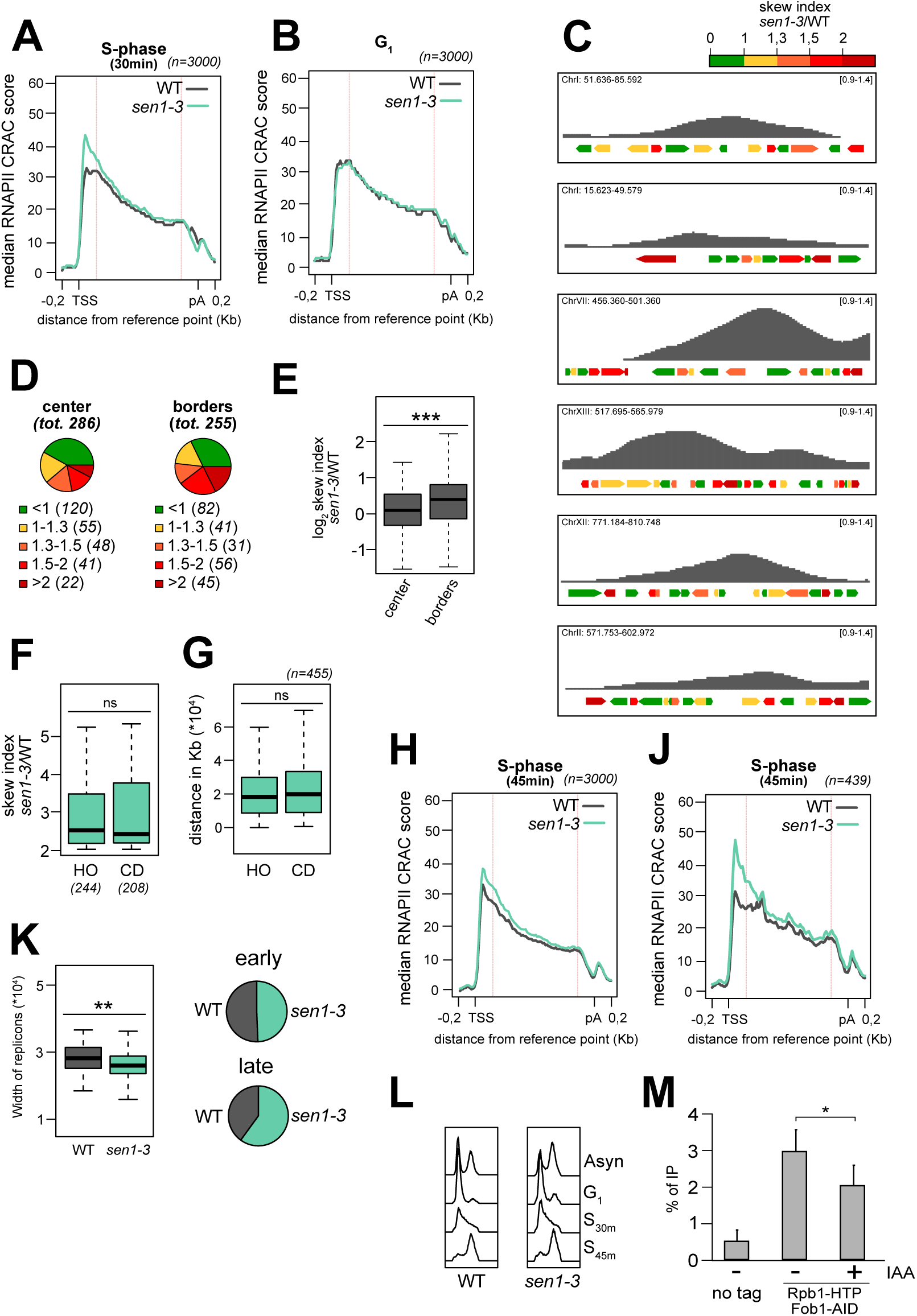
**A)** and **B)** As in Figure 2A, but on cells synchronously released in S-phase and collected 30 min after replication onset or arrested in G_1_, respectively. The 3000 genes with the highest transcription levels have been used. **C)** As in Figure 2F, additional representative snapshots illustrating the frequent co-localisation of the replication forks (minima of the DNA copy number signal) and genes displaying a 5’-end skewed RNAPII distribution pattern in *sen1-3* cells (coloured in red). **D)** Pies indicating the total number of genes belonging to the high or low skew ratio classes as in C, overlapping the ‘borders’ or ‘center’ of replicons. **E)** Boxplot comparison of the skew index ratio (*sen1-3*/WT) of RNAPII CRAC signal for genes included within 7.5 Kb of replicons borders or in their center. **F)** Boxplot comparison of the skew index ratios (*sen1-3*/WT) of RNAPII signal for genes oriented in head-on (HO) or in co-direction (CD) relative to the position on the nearest activated ARS as detected by DNA copy number analysis. Genes in both groups have a replication-dependent accumulation of RNAPII in their 5’-end. **G)** Distribution of distances for each affected gene from the closest HO and CD origins. Affected genes are not preferentially replicated in one configuration over the other. **H)** Meta-analyses as in Figure 2A, but on cells synchronously released in S-phase and collected 45 min after replication onset. **J)** As in Figure 2C, but on cells synchronously released in S-phase and collected 45 min after replication onset. The analysis was performed on the subset of genes affected at this time point. **K)** Left: comparison of the width of the replicons detected by DNA copy number analysis from WT and *sen1-3* cells synchronously grown in S-phase and collected 30 min after replication onset. Right: pies indicating the total number of activated origins that were retrieved from DNA copy number analysis in wild-type (WT) and *sen1-3* cells. Origins were divided in early and late according to their replication timing in wild-type cells. **L)** Examples of cell cycle analysis by flow cytometry from the cells used for the experiments shown in Figure 2 and S2. **M)** RNAPII (Rbp1) levels by ChIP-qPCR at the rRFB upon conditional depletion of Fob1 via the auxin degron system. The average level of 3 independent experiments is plotted. In all panels: *p<0.05, **p<0.01, ***p<0.001.

**Supplementary Figure 3 related to Figure 4:**
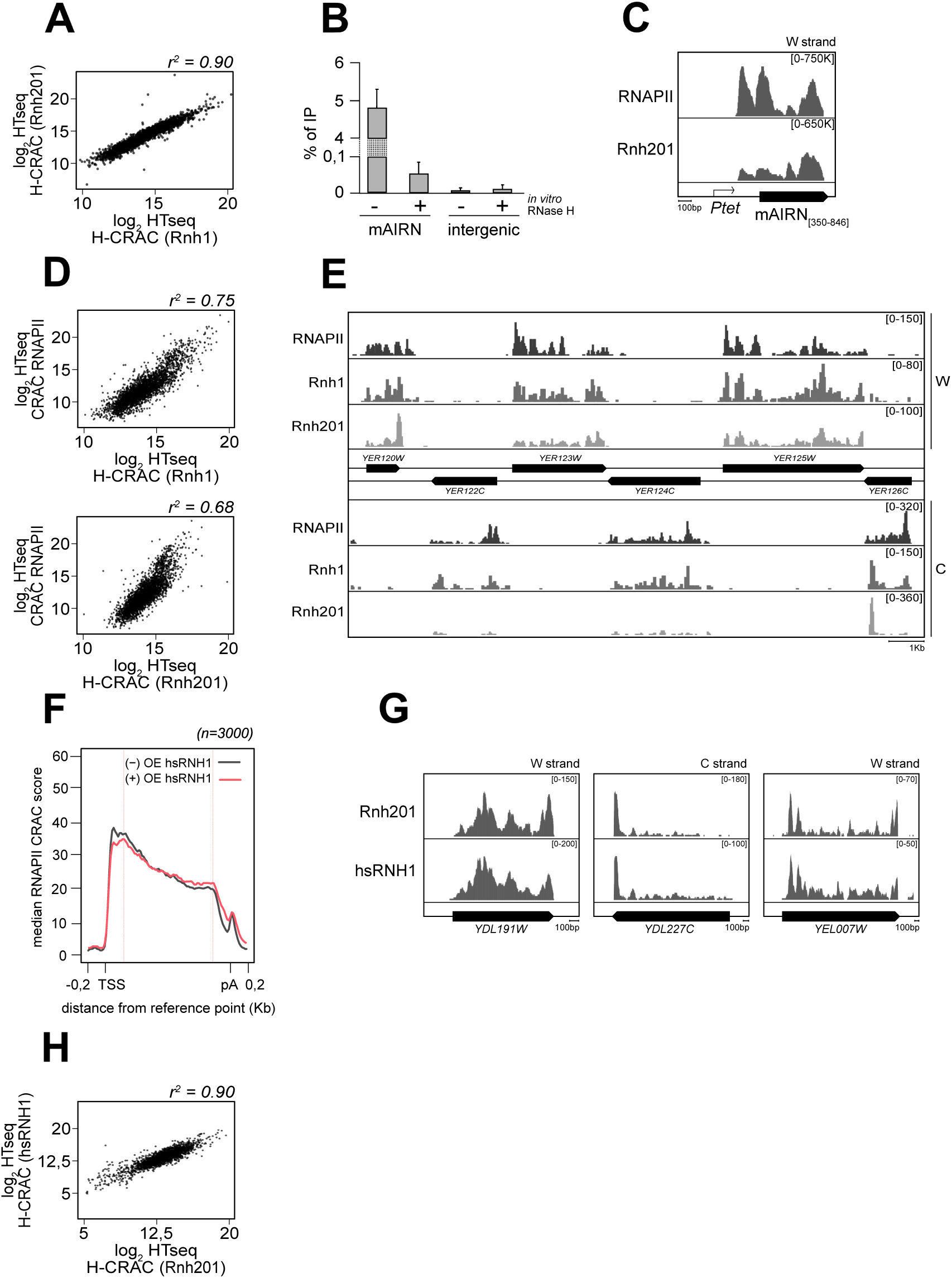
**A)** Dispersion plot of the log_2_ values obtained from Rnh1 H-CRAC and Rnh201 H-CRAC for the 3000 most transcribed genes in asynchronous cells. The coefficient of determination (r^2^) is shown. **B)** DNA:RNA Immunoprecipitation followed by quantitative PCR (DRIP-qPCR) from cells transformed with a plasmid carrying the mAIRN sequence expressed under control of the *pTet*promoter as indicated on the scheme in panel C (only the R-loop-forming region corresponding to the 350-848 nt interval of the mouse gene was cloned). The percentage of immunoprecipitated material is plotted on y-axis. The DRIP signal from an intergenic region located nearby to the *HO* locus was used as a negative control. Samples were treated or not with RNase H *in vitro* prior to immunoprecipitation as indicated. **C)** Read coverage for RNAPII (CRAC) and Rnh201 (H-CRAC) on the plasmid-borne mAIRN sequence. **D)** Dispersion plots illustrating the correlation between transcription (RNAPII CRAC) and R-loop levels, as determined by H-CRAC (top: Rnh1; bottom: Rnh201). **E)** Integrative Genomics Viewer (IGV) representative screenshot of a chromosomal region illustrating the marked directionality of H-CRAC signals for both Rnh1 and Rnh201 as indicated. **F)** Metagene analyses of RNAPII CRAC signal at coding genes aligned on their TSS and on their pA site in WT cells transformed with a plasmid overexpressing hsRNH1 (+) or an empty plasmid (-). Genes are only scaled in between the red lines. **G)** Representative snapshots illustrating the similarities between the H-CRAC signal obtained with yeast Rnh201 or human RNH1 (hsRNH1). **H)** Dispersion plot as A but comparing Rnh201 to hsRNH1.

**Supplementary Figure 4 related to Figure 4:**
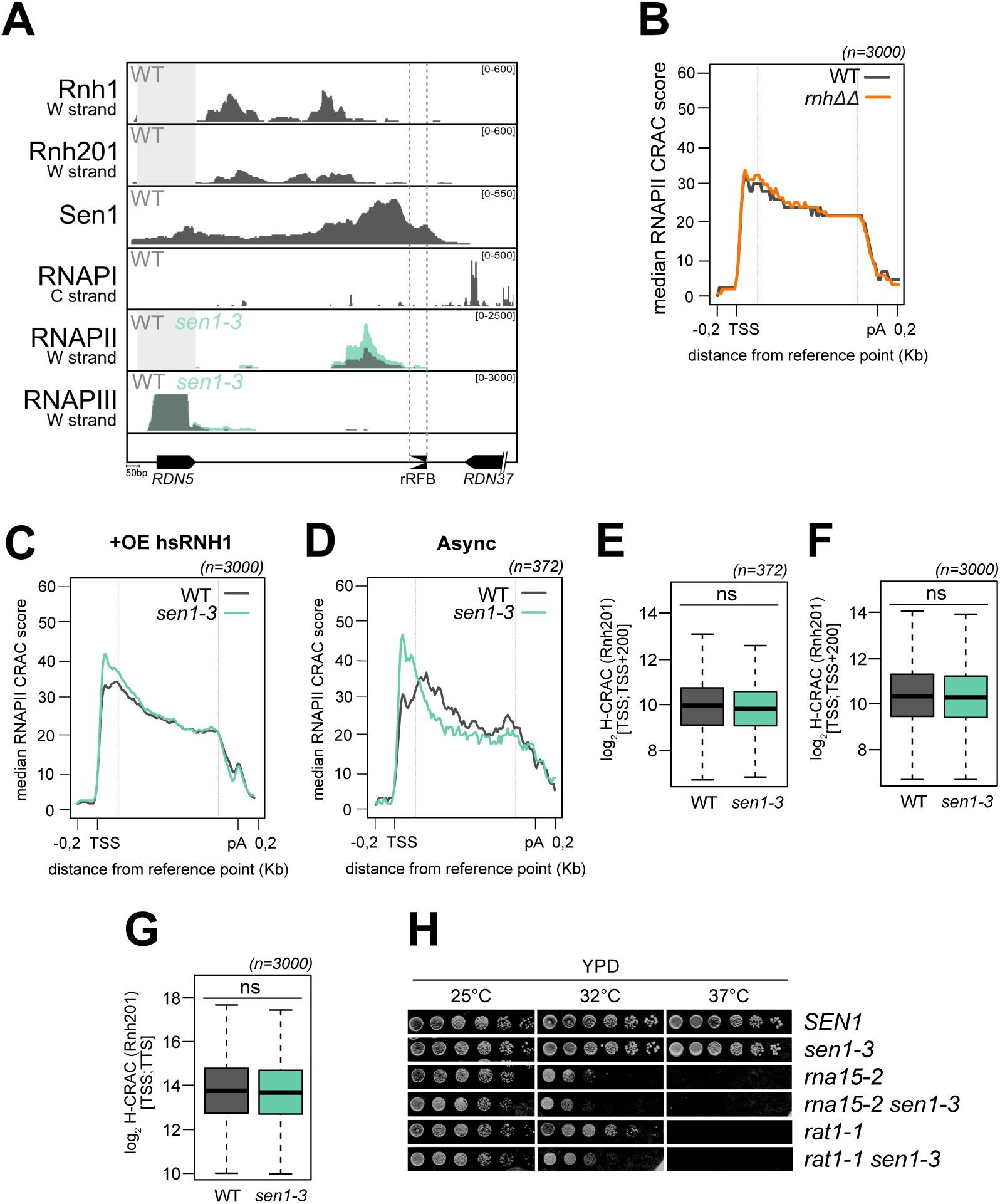
**A)** Distribution of CRAC signals of the indicated proteins at the rRFB. Data for RNAPI (Turowski et al., 2020) and RNAPIII (Xie et al., 2021) indicate that accumulation of RNAPII is unlikely due to conflicts with RNAPI or RNAPIII. Sen1 ChIP-exo signals (Rossi et al., 2021) demonstrate the presence of Sen1 at this site of TRCs. The strand shown is indicated for each protein, with the exception of Sen1 because ChIP data are not directional. **B)** Metagene analyses of RNAPII CRAC signal at coding genes aligned on their TSS and pA site in *rnh1Δ rnh201Δ* (*rnhΔΔ*) or WT cells. Genes are scaled only in between the red lines. **C)**As in B but for WT and *sen1-3* cells transformed with a plasmid over-expressing human RNH1. Note that the 5’-end RNAPII accumulation observed in *sen1-3* cells is not lost in these conditions. **D)** As in Figure 2C but on asynchronously dividing cells and on the subset of genes affected in this condition. **E)** and **F)** Boxplot comparisons of the H-CRAC signal on the interval [TSS; TSS+200] for the same group of genes shown in Figure S4D or for the most transcribed 3000 genes, respectively. **G)** As in F, but for the signal along the full gene [TSS; TTS]. **H)** Growth assay of *rat1-1 sen1-3* and *rna15-2 sen1-3* cells compared to single mutants. Serial dilutions of the indicated strains were incubated for 3 days at the indicated temperature. Growth was performed on the same plates. The synthetic effect of the combination of the *rat1-1* and *sen1-3* alleles is unlikely due to defective termination of mRNA-coding genes (due to the *rat1-1* mutation), because the even stronger termination defect induced by mutation of Rna15, does not result in a similar synthetic phenotype when associated to *sen1-3*.

**Supplementary Figure 5 related to Figure 5:**
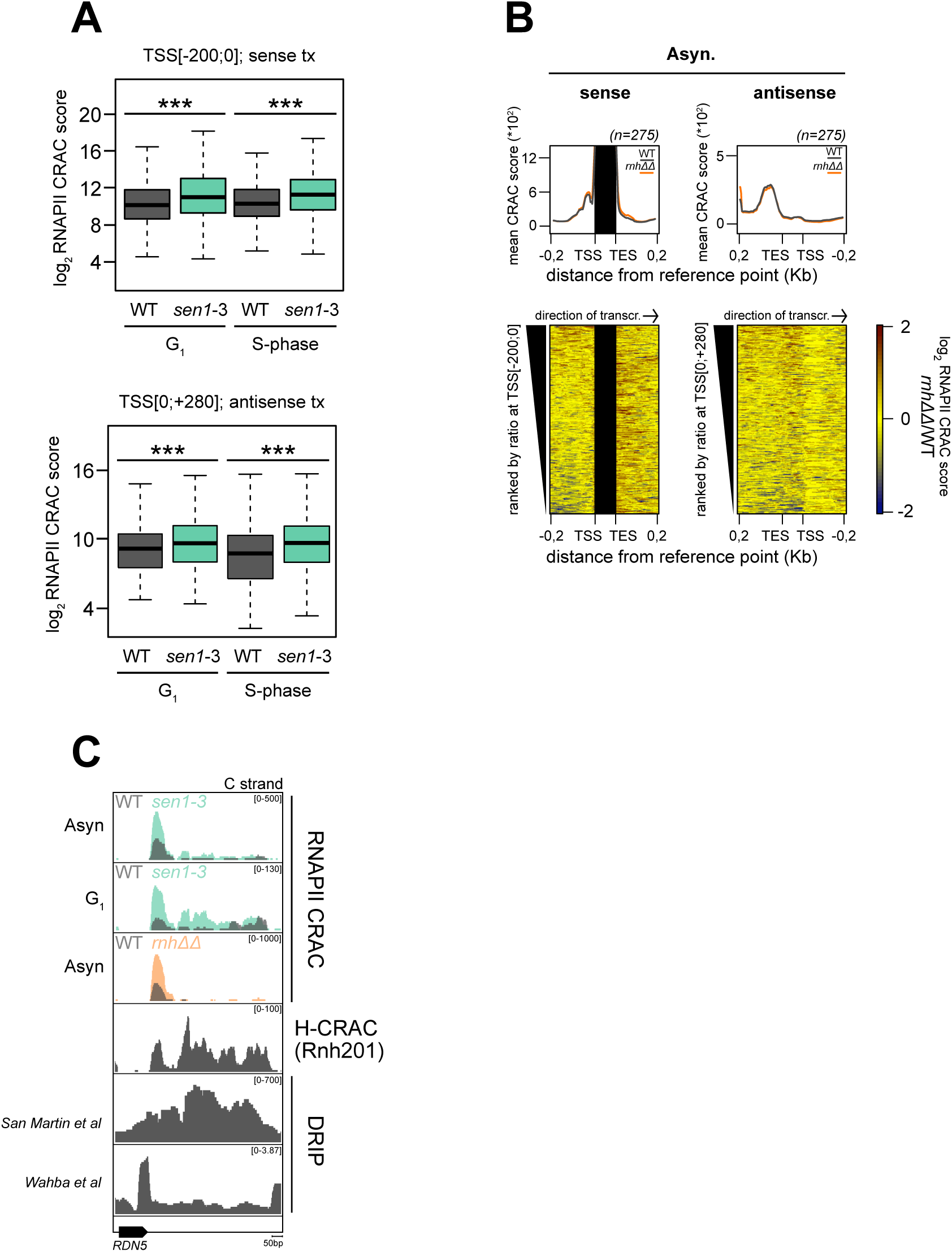
**A)** Boxplot comparison of sense (top) or antisense (bottom) transcription (RNAPII CRAC) at tRNA genes in WT and *sen1-3* cells, arrested in G_1_ or synchronised in S-phase (30 min) as indicated. *** p-value<0.001. **B)** Heatmap analyses as in Figure 5B but comparing RNases H deleted cells to a WT condition. **C)** Accumulation of RNAPII antisense to the *RDN5* gene in WT, *sen1-3* and *rnh1Δ rnh201Δ* cells as in Figure 5A for tRNAs. R-loops levels from H-CRAC and DRIP-seq (San Martin-Alonso et al., 2021; Wahba et al., 2016) are also shown.

**Supplementary Figure 6 related to Figure 6:**
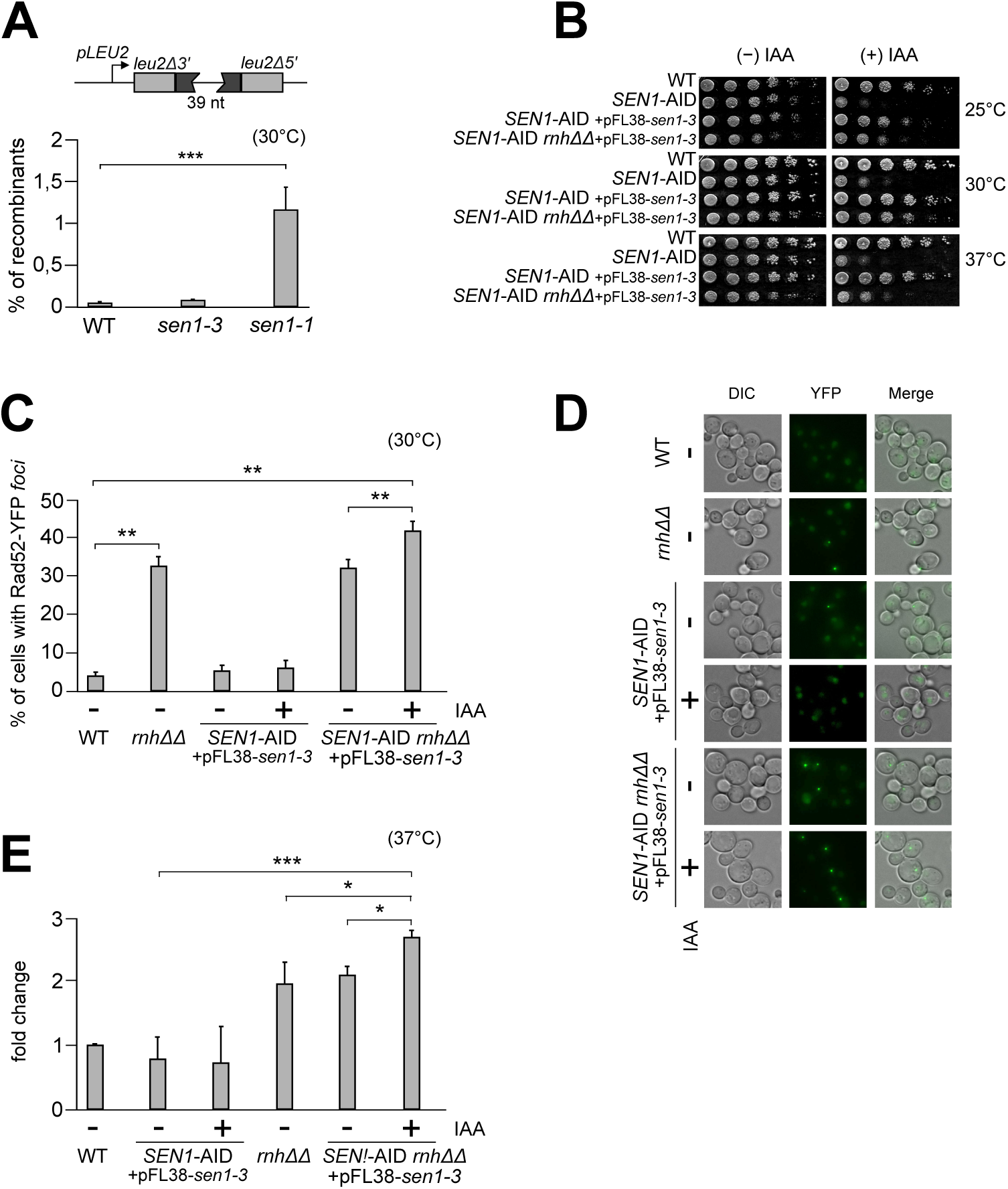
**A)** Frequency of transcription-associated recombination (TAR) events assessed using the pL (Prado and Aguilera, 2005) reporter in WT, *sen1-3* and *sen1-1* cells. The reporter plasmid (schematised above the graph) contains the *LEU2* gene interrupted by a 39 nt insertion flanked by homology repeats. Transcription activation in the absence of leucine induces damage and recombination between the two repeats, which reconstitutes a functional *LEU2* gene. **B)** Growth assay of inducible triple mutant used for the analyses shown in Figures 6 and S6. Note that the establishment of the phenotype is partial, possibly due to the partial depletion of Sen1 or a suppression effect of slight Sen1-3 overexpression. **C)** As in Figure 6B, but growth was performed at 30°C. **D)** Representative examples of the microscopy data shown in Figure S6C. **E)** Quantification of H2A Ser129 phosphorylation detected by western blot in Figure 6C. Cells were grown in logarithmic phase at 30°C, and then shifted at 37°C for 1 hour. The fold change relative to the WT levels is shown for the indicated conditions. Pab1, Nrd1 and Nab3 were used as loading controls. Error bars represent standard deviations. For all panels: *p<0.05, **p<0.1, ***p<0.001.

**Supplementary Figure 7:**
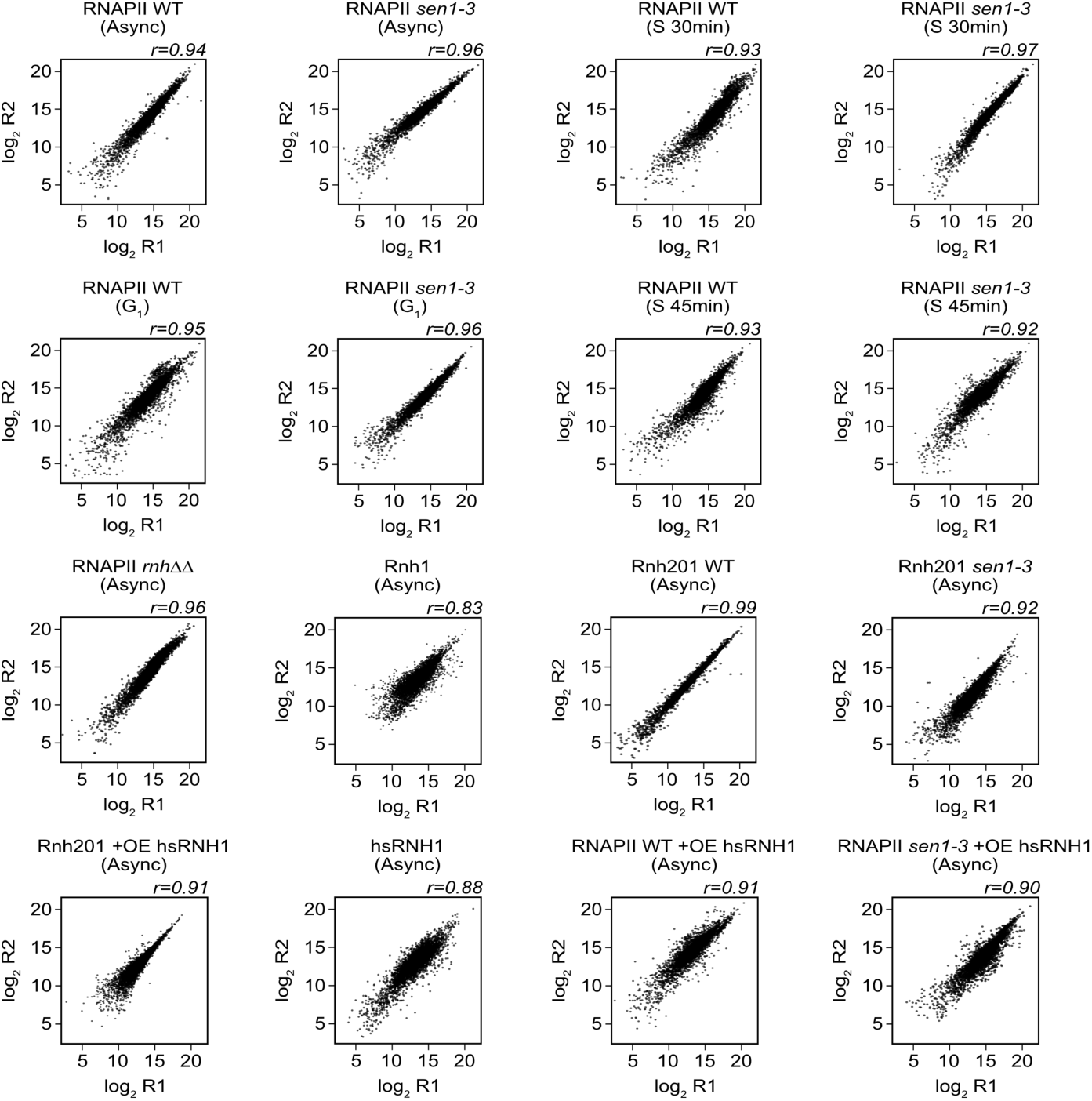
Correlation plots between replicates of the CRAC experiments shown in this study. The Pearson correlation score (r) is shown at the top right of each plot.

## Notes

### Competing Interest Statement

The authors have declared no competing interest.

